# Apolar mode of gastrulation leads to the formation of polarized larva in a marine hydroid, *Dynamena pumila*

**DOI:** 10.1101/2021.02.25.432844

**Authors:** Alexandra A. Vetrova, Tatiana S. Lebedeva, Aleena A. Saidova, Daria M. Kupaeva, Yulia A. Kraus, Stanislav V. Kremnyov

**Affiliations:** Laboratory of Morphogenesis Evolution, Koltzov Institute of Developmental biology RAS, Moscow, Russia; Department of Embryology, Faculty of Biology, Lomonosov Moscow State University, Moscow, Russia; Department for Molecular Evolution and Development, Centre of Organismal Systems Biology, University of Vienna, Vienna, Austria; Department of Cell Biology and Histology, Faculty of Biology, Lomonosov Moscow State University, Moscow, Russia; Department of Evolutionary Biology, Faculty of Biology, Lomonosov Moscow State University, Moscow, Russia

**Keywords:** cnidaria, morphogenesis, gastrulation, cell migration, evo-devo

## Abstract

**Background:** In almost all metazoans examined to this respect, the axial patterning system based on canonical Wnt (cWnt) signaling operates throughout the course of development. In most metazoans, gastrulation is polar, and embryos develop morphological landmarks of axial polarity, such as blastopore under control/regulation from Wnt signaling. However, in many cnidarian species, gastrulation is morphologically apolar. The question remains whether сWnt signaling providing the establishment of a body axis controls morphogenetic processes involved in apolar gastrulation.

**Results:** In this study, we focused on the embryonic development of *Dynamena pumila*, a cnidarian species with apolar gastrulation. We thoroughly described cell behavior, proliferation, and ultrastructure and examined axial patterning in the embryos of this species. We revealed that the first signs of morphological polarity appear only after the end of gastrulation, while molecular prepatterning of the embryo does exist during gastrulation. We have shown experimentally that in *D. pumila,* the morphological axis is highly robust against perturbations in cWnt activity.

**Conclusion:** Our results suggest that morphogenetic processes are uncoupled from molecular axial patterning during gastrulation in *D. pumila*. Investigation of *D. pumila* might significantly expand our understanding of the ways in which morphological polarization and axial molecular patterning are linked in Metazoa.

## Introduction

Canonical Wnt (cWnt) signaling controls the establishment of body axis polarity during embryonic development in most metazoans (Petersen, Reddien, 2009). Axial molecular patterning relies on a gradient distribution of cWnt signaling components. When gastrulation morphogenetic movements are linked to the axial patterning and occur at a specific position to the embryonic axes (Leclère et al., 2016), they are referred to as polar. Most metazoan species examined in this respect exhibit one of the modes of polar gastrulation, with invagination as a prime example (Kominami, Takata, 2004; Magie et al., 2007; Winkley et al., 2020). In these cases, embryo morphology is coupled with axial molecular patterning throughout gastrulation.

In species of the phylum Cnidaria, placed as a sister group to Bilateria, cWnt signaling patterns the primary oral-aboral body axis, controls oral identity during larval body plan formation, and coincides with a single region of gastrulation morphogenesis in the case of polar gastrulation (e.g., *Clytia, Nematostella*) (Lee et al., 2007; Momose et al., 2008). However, in species within the class Hydrozoa (Cnidaria), one can find apolar gastrulation as well.

Apolar gastrulation occurs as multipolar ingression of individual cells or delamination (Martin et al., 1997, Kraus, Markov, 2017). One can distinguish between two types of delamination (Metschnikoff, 1886). During primary delamination, external cells divide perpendicularly to the surface of the embryo. As a result, one of the cells remains in the outer layer, and the other goes inside, where it joins the presumptive endoderm. During secondary delamination, external cells start to epithelialize throughout the embryo until they segregate themselves from the inner cell mass completely. These external cells form an ectodermal epithelial layer while inner cell mass forms an endoderm.

In apolar gastrulation, there are no morphological landmarks, which indicate an embryonic polarity. For instance, *Hydractinia echinata* and *Gonothyrae loveni* gastrulate in an apolar fashion, and the direction of the oral-aboral axis becomes evident only at the end of gastrulation (Kraus et al., 2014; Burmistrova et al., 2018). The molecular basis of axis formation within species with apolar gastrulation is not explored sufficiently. Expression patterns of cWnt signaling components were demonstrated in *H. echinata* only for stages with clear morphological polarization (Plickert et al., 2006; Hensel et al., 2014). However, it remains unclear how cWnt signaling establishes the oral region in these species and whether morphogenetic processes colocalize with domains of cWnt component expression during morphologically apolar stages of embryonic development. One can consider several possibilities. The first option is that a single oral region of cWnt activity exists in embryos, but gastrulation morphogenetic processes are uncoupled from it. The second option is that morphogenetic processes of apolar gastrulation are coupled with multiple regions of cWnt signaling throughout an embryo when a single oral domain is established later. In such a case, it might resemble the situation during *Nematostella* embryonic aggregate development (Kirillova et al., 2018).

To answer these questions, we have carefully studied the embryonic development of *Dynamena pumila*, a species known for an extremely variable morphology during apolar gastrulation (Kraus, Cherdantsev, 1999). *D. pumila* is a widespread colonial hydroid, which inhabits the littoral zone. Сhitinous theca cover tissues of the colony (Figure 1A). Gamete maturation takes place in strongly reduced medusae, which develop inside the so-called gonothecae (Figure 1B, C) (Teissier, 1922). Unlike spawning species, oocytes do not emerge into the water column. They shed into the acellular mucous acrocyst, which everts from the gonotheca and remains attached to a colony (Figure 1D).

**Figure 1.**
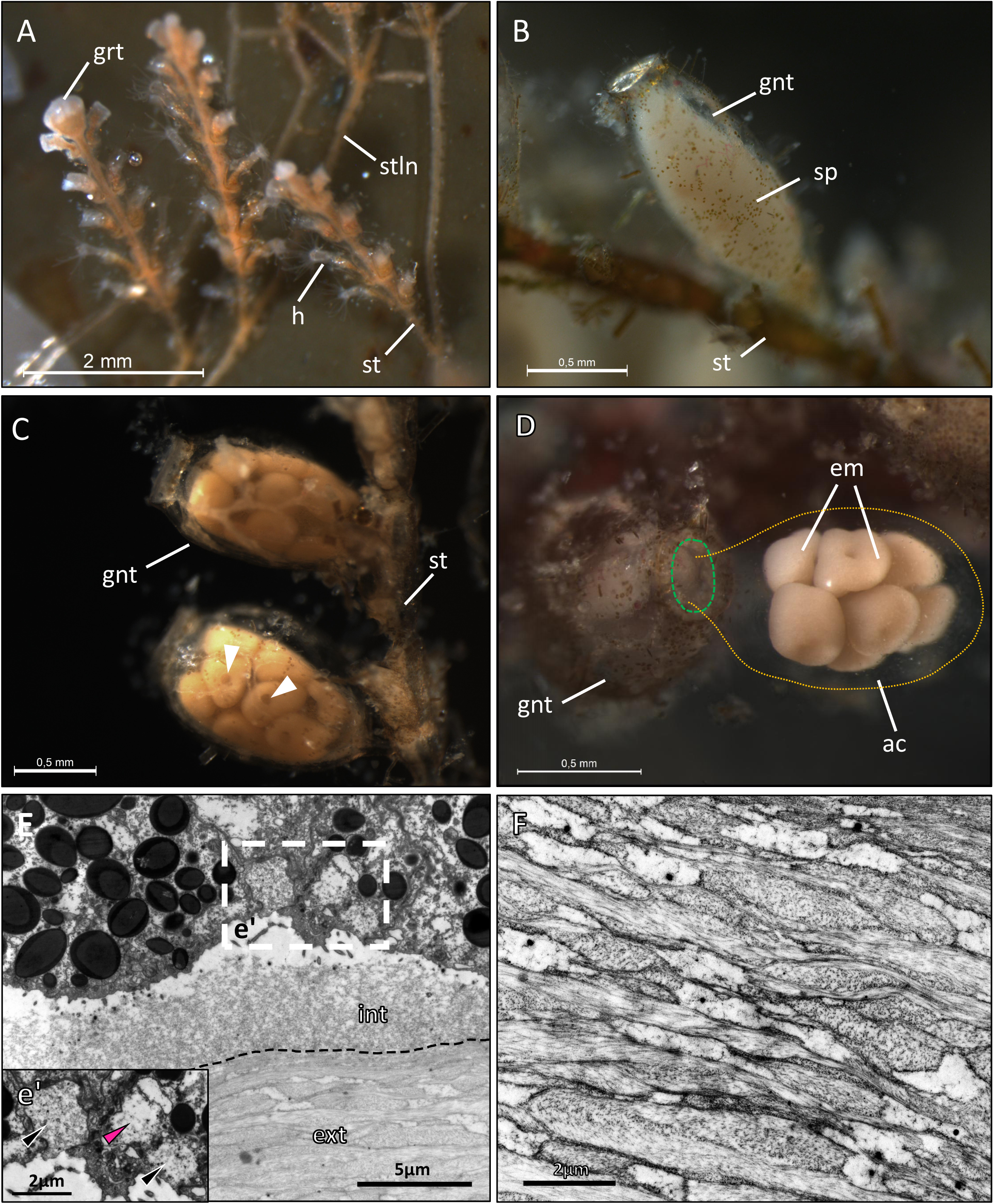
(A) The colony of *Dynamena pumila*. grt -growing tip; h - hydrant; st - stem; stln - stolon. (B) Male and (C) female gametes maturation. Gonotheca (gnt) surrounds sperm (sp) or oocytes. Arrowheads indicate nuclei of oocytes. (D) Embryos (em) at the gastrula stage in mucous acrocyst (ac, yellow dotted line), coming out from the opening (green dotted line) of gonotheca. (E) The mucus of an acrocyst: irregular internal (int) and structured external (ext) layers. The mucus layer boundary is shown by the dotted line. (e’) Mucus-containing vesicles (pink arrowhead) in embryonic cells. Mucus extrusion from vesicles (black arrowheads). (F) The structure of acrocyst’s external layer. TEM: (E-F)

The mucus of the acrocyst is noticeably structured and several morphologically distinguishable zones can be observed (Figure 1E). The mucus, likely, is produced by colony tissues. The embryos may contribute to mucus production. Large mucus-containing vesicles at the different stages of exocytosis can be observed in outer cells (Figure 1e’). The external mucus is composed of many interwoven strands (Figure 1F). Most likely, such an organization of mucus gives strength to the acrocyst.

Inside the acrocyst, embryonic development lasts till the formation of the motile planula larva. After planula formation, it leaves the acrocyst and settles to give rise to a new colony after metamorphosis (Bagaeva et al., 2019).

The first description of *D. pumila* development was performed by Teissier (1923). Since then, *D. pumila* regularly comes under scrutiny: embryonic development and molecular aspects of colony formation were previously investigated (Kraus, Cherdantsev, 2003; Kraus, 2006; Bagaeva et al., 2019). However, embryonic cell ultrastructure, the molecular basis of morphogenesis, and body axis formation have not been explored in sufficient detail throughout the development of *D. pumila*.

In this study, we provide a detailed description of *D. pumila* embryonic development using in vivo observations (Figure 2), laser confocal, light, and electron microscopy. Since *D. pumila* is a littoral species and withstands varying temperatures in natural conditions, we examined its development in the temperature range between +12-16° C. We thoroughly characterized embryonic morphology, ultrastructure, and cell behavior up to the formation of planula larva and examined the molecular mechanisms of axial patterning during the development of *D. pumila*. Our findings enhance understanding of the morphogenetic mechanisms underlying the peculiar gastrulation of *D. pumila* and reveal the molecular basis of axis patterning in this species.

**Figure 2.**
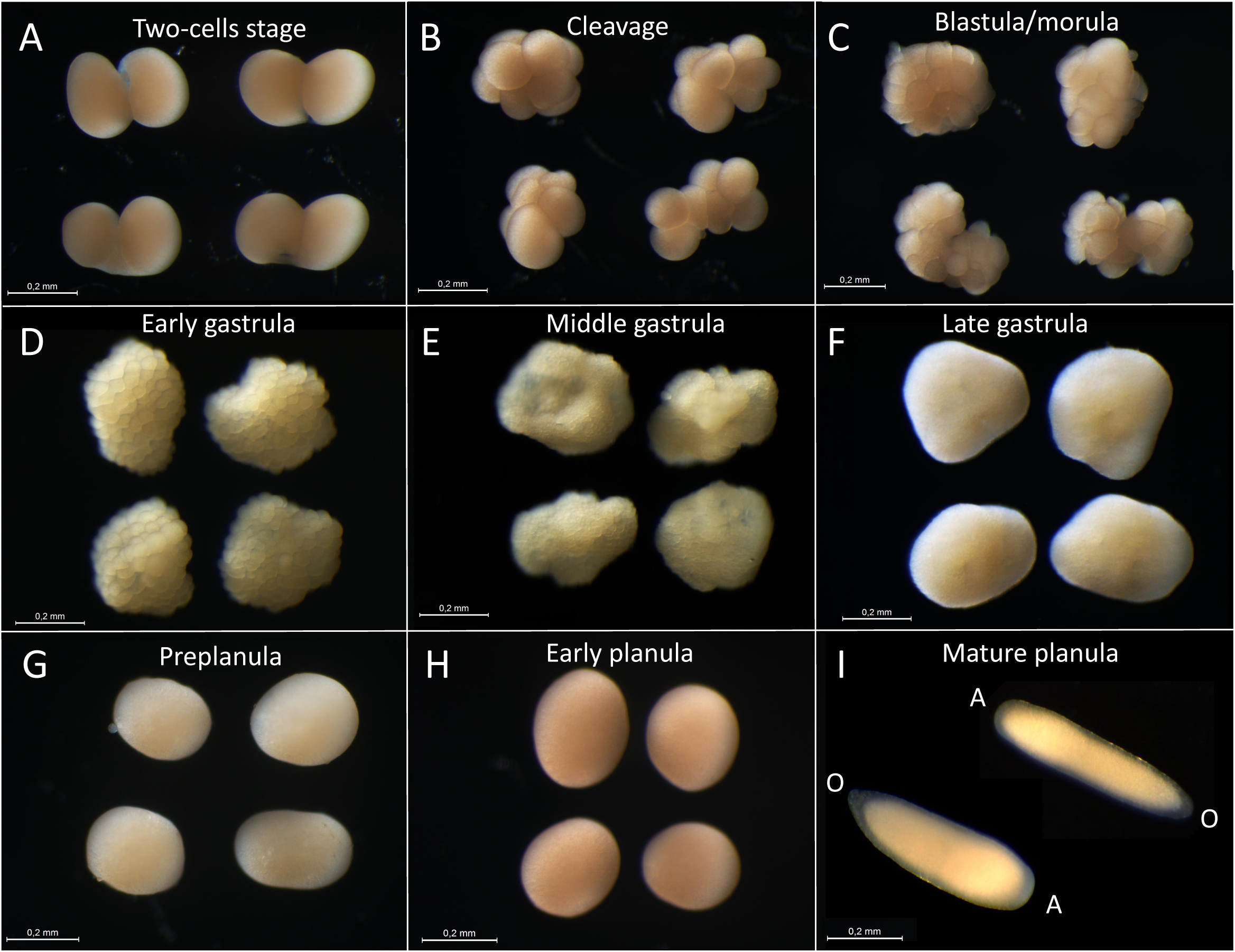
Embryonic development of *D. pumila*. A - aboral pole; O - oral pole. Scale bar: 0.2 mm

## Results

### Cleavage

Zygotes are oval or almost spherical in shape and about 230-350 μm in diameter (Figure 3A). The first cleavage furrow originates unilaterally on the animal pole of a zygote from two sites of intrusion (Figure 3B: arrows). It seems that these intrusions elongate along the surface of the embryo towards each other and merge into a channel. Further, the channel breaks (Figure 3C), leaving its contracted remnants as crests on both sides of the formed furrow (Figure 3D).

**Figure 3.**
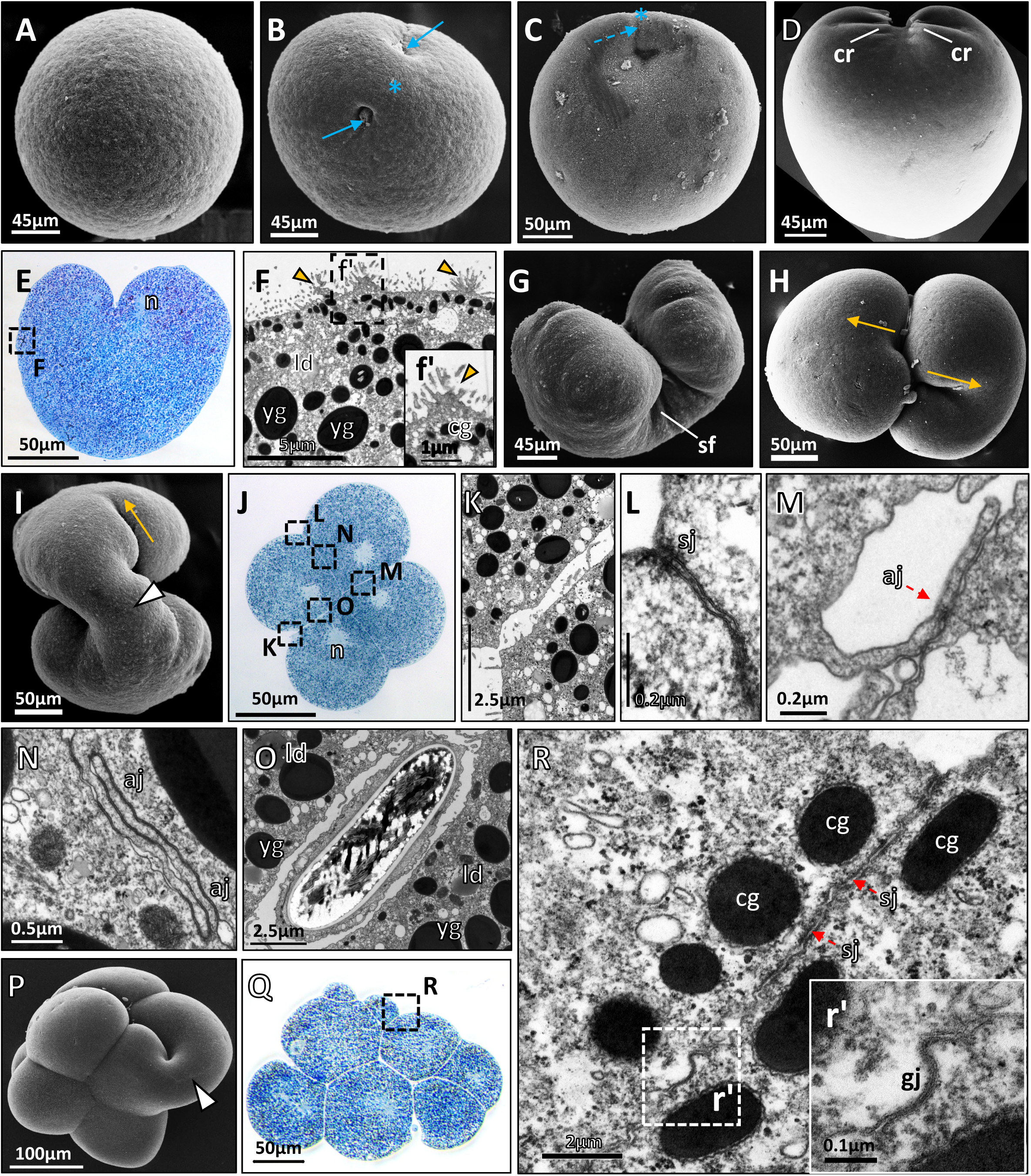
Cleavage. (A) Zygote. (B, C) Start of cleavage. Blue asterisks mark the animal pole. Blue solid arrows point to sites of intrusion, blue dotted arrow - to the site of future break. (D) Break of the cleavage furrow. cr - membrane crest. (E) First cleavage. n - nucleus. (F, f’) Cortical cytoplasm: cg - cortical granules; ld - lipid droplet; vc - vacuole; yg - yolk granule. Yellow arrowheads point to the bundle of microvilli. (G) Embryo with a deep cleavage furrow. sf - surface folds. (H, I) Start of the second cleavage: view from the animal pole - H, from the vegetal pole - I. Yellow arrows show the direction of furrow elongation. White arrowhead points to the cytoplasmic bridge between blastomeres. (J) 4-cells stage. (K) Adjacent surfaces of blastomeres. (L) Septate junction (sj) in the subapical region. (M) Spot-like adherens junction (aj, red arrow). (N) Adherens junctions. (O) Maternal nematocyte between blastomeres. (P) Non-synchronous cleavage. (Q) Central section of cleavage-stage embryo. (R) Apico-basal distribution of cell junctions. (r’) Gap junction (gj). SEM: (A-D, G-H, P). Semi-thin sections: (E, J, Q). TEM: (F, f’, K-O, R, r’).

The cleavage furrow elongates along the animal-vegetative axis. Its direction can be identified by nuclei forming near the animal pole (Figure 3E). Embryo cytoplasm is enriched with lipid droplets and big spherical electron-dense granules, which were previously identified as yolk granules in embryos of other hydrozoan species (Kraus et al., 2014; Burmistrova et al., 2018). Elongated electron-dense granules 0.4-0.9 μm in length are distributed between yolk granules and in the cortical cytoplasm. Similar granules were described in embryos of *Pennaria tiarella* and *H. echinata*, but their nature is still unknown (Martin, Thomas, 1977; Kraus et al., 2014). Gathered in bundles, microvilli cover the embryo surface (Figure 3F, f’: arrowheads). As the elongation of cleavage furrow proceeds, the embryonic surface folds at its bottom (Figure 3G).

The furrows of the second cleavage originate unilaterally on the contact surfaces of blastomeres (Figure 3H). The second round of cleavage may start before the completion of the first one (Figure 3I). Displacement of blastomeres occurs concurrently with cleavage furrow formation (Figure 3H, I).

At the 4-cells stage, nuclei shift towards the center of the embryo (Figure 3J). Blastomeres are packed quite loosely (Figure 3K), but cell junctions start to form already. It is possible to distinguish septate junctions (Figure 3L) closer to the outer surface of the blastomere and presumably adherens junctions lying deeper. Spot-like adherens junctions start forming between the filopodia of one blastomere and the body of the neighboring blastomere (Figure 3M). Then the contact area increases and ties the blastomeres tightly (Figure 3N).

During cleavage and at the later stages, nematocytes can be found between blastomeres (Figure 3O). It is supposed that these cells are derived from the colony tissues. It was shown that during acrocyst formation, maternal nematocytes fall into the mucus surrounding the eggs and infiltrate the space between embryonic cells later (Teissier, 1923).

After the second cycle, cleavage loses its synchrony but remains unilateral. The cleavage furrow originates at the center of the embryo and extends towards its surface (Figure 3P). Embryos differ in cell packing highly due to active cell displacement and the varying rate of cytokinesis (Figure 2B; 3P). Blastomeres are packed tightly and the blastocoel is absent (Figure 3Q). Thus, the embryo loses morphological polarity at the cleavage stage.

Cell junction formation continues at the 8-cell stage. Septate junctions up to hundreds of nanometres in length are detectable in the subapical blastomere regions, presumptive gap junctions are located more basally (Figure 3R, r’).

Cleavage lasts 3-4 hours from the initiation of the first furrow until the 16-32-cell stage at +12-18^о^C.

### Blastula/morula

Embryos at the 16-32-cell stage may become a blastula with a small blastocoel (Figure 4A), or a morula if the blastocoel is absent, since several cells reside within the embryo (Figure 4B). It seems that intermediate forms are also possible. This stage lasts only for 1-2 hours.

**Figure 4.**
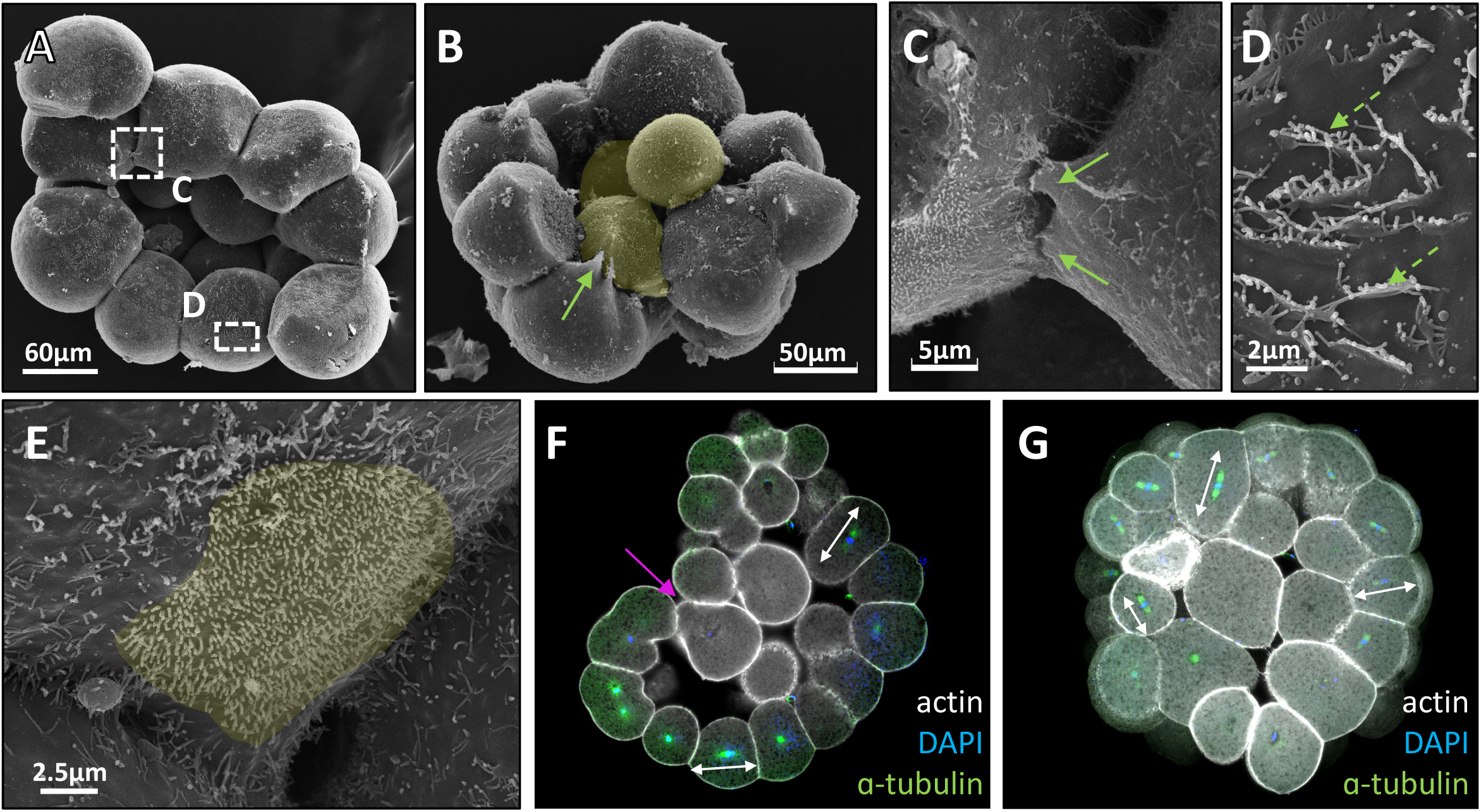
Transient blastula/Morula. (A) Transient blastula split into halves. (B) Morula split into halves. Inner cells are artificially colored in yellow. Solid arrow points to lamella-like protrusions. (C) Lamella-like (green solid arrows) and (D) ridge-like (green dotted arrows) protrusions on the lateral cell surfaces. (E) Microvilli on a non-contact part of the basal cell surface are artificially colored in yellow. (F, G) Morula-type embryos, CLM central optical section. White double-arrows show the direction of mitotic spindles. Magenta arrow points to the cell, presumably shifting inside. SEM: (A-E).

The embryo is composed of roundish and slightly triangular cells with rounded external surfaces. Their internal-lateral surfaces form large lamellipodia contacting neighboring cells (Figure 4B, C: arrows). They form multiple cytoplasmic membrane ridges on the adjacent surfaces of blastomeres in the zones of intercellular contact (Figure 4D: arrows). These ridges can be found during the early stages of development only and disappear before gastrulation.

Cell surfaces are covered with evenly distributed minute microvilli in non-contact zones, including the internal cell surfaces (Figure 4E). Microvilli are highly uncharacteristic features for basal cell surfaces in general.

From the 64-cell stage approximately, there are only morula-type embryos of irregular shape (Figure 4F, G), while blastula-type embryos were not observed. Two options exist for how cells might get inside and fill the blastocoel cavity. Some mitotic spindles are orientated perpendicularly to the surface of the embryo (Figure 4F, G: double arrows), so one of the daughter cells goes inside. Another option is a rearrangement of cells (Figure 4F: arrow).

### Early gastrula

From the 128-cell stage on, epithelial sheet fragments composed of external cells can be observed in the embryo (Picture 5A, B). An embryonic cell sheet is considered epithelial if the cells are clearly polarized and aligned, and have elaborate intercellular junctions (Knust, Bossinger, 2002). We consider this stage as the start of gastrulation. Overall, gastrulation lasts for about 40 hours.

Prior to gastrulation, the embryonic surface consists of flat and curved regions in such a way that flat surfaces of the embryo are surrounded by curved surfaces (Figure 5A). Epithelization starts, primarily, on the curvature of the embryonic surface simultaneously in several regions (Picture 5A, B). A scheme of morphogenesis of local epithelial sheet fragments was proposed previously (Kraus, 2006). In sites of epithelization, the cell shape changes from roundish to pyramidal or columnar with a slightly widened apical domain. It seems that not only external cells are involved in this process: we could observe curling «rosettes» at the free edges of epithelial sheet fragments (Figure 5A). With the start of the epithelization process, the variability in the structure of embryos increases significantly.

**Figure 5.**
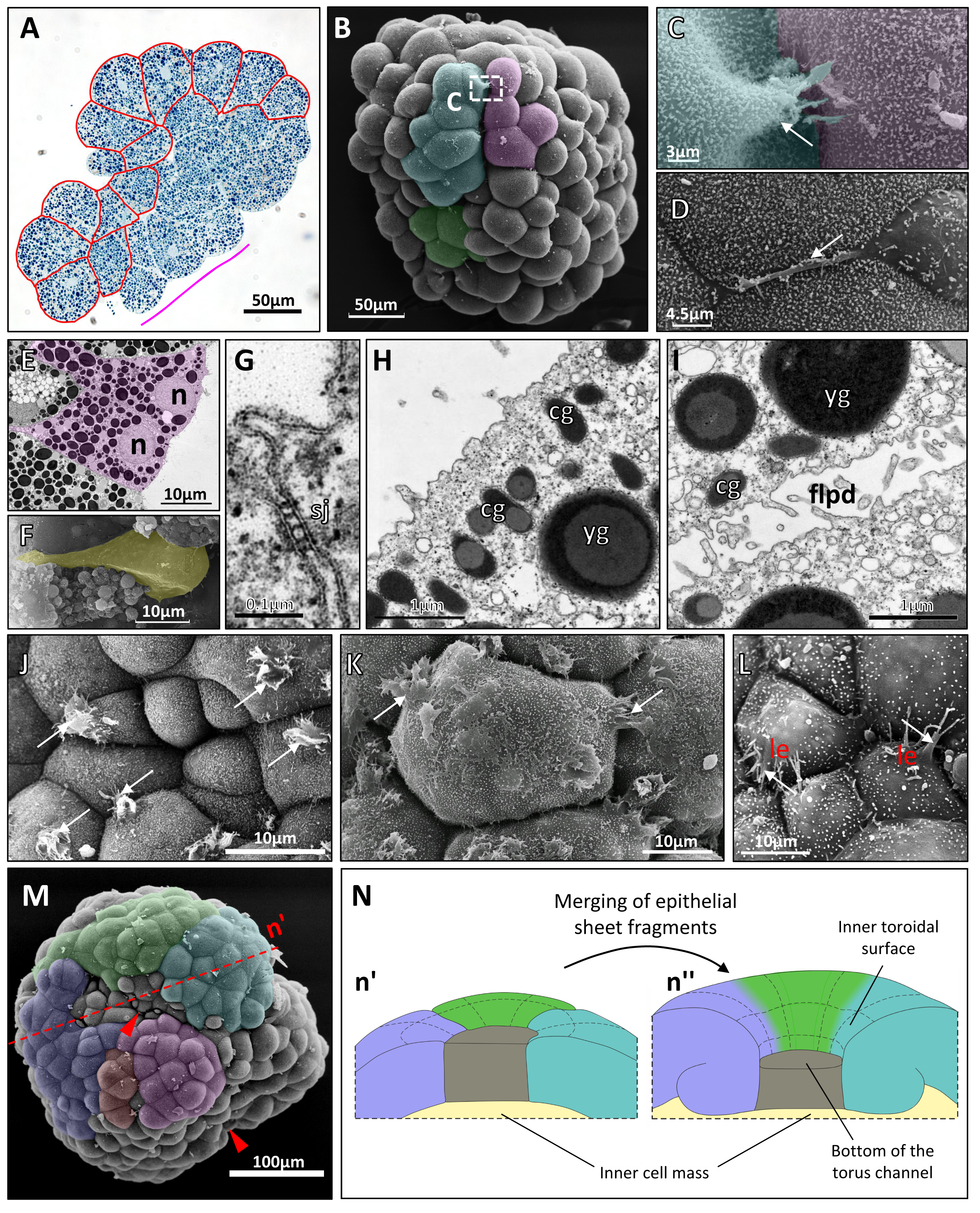
Early gastrula stage. (А) Semi-thin section. Red lines accentuate the shape of cells in epithelial sheet fragments (esf). The magenta line shows the flat region of the embryonic surface. (B) Esfs are colored artificially. (C) Lamellopodia (arrow). (D) Filopodium (arrow). (E) Nuclei (n) are shifted to the apical domain of esf cells. Cells are highlighted in magenta. (F) Outer cell with constricted apex is colored in yellow. (G) Septate junction (sj) in the sub-apical region of outer cells. (H) The apical domain of the esf cell. cg - cortical granule; yg - yolk granule. (I) Adjacent surfaces of inner cells. flpd - filopodia. (J-L) Stages of the formation of cell protrusions (arrows). le - leading edge. (M) Merging esfs are colored artificially. Red arrowheads indicate forming torus openings. The red dotted line represents a cut through the embryo, schematically shown on (n’). (N) Scheme of toroidal surface formation: n’ - early gastrula stage, n” - middle gastrula stage. SEM: B-D, F, J-M. TEM: E, G-I.

We detected cells with filopodia and lamellipodia in developing epithelial sheet fragments (Figure 5C, D: arrows), which seem to establish contacts with cells of other epithelial sheet fragments (figure 5B, C).

Cells mostly attain a triangular or columnar shape in well-developed epithelial sheet fragments (Figure 5E). Some external cells between epithelial sheet fragments have reduced apical and expanded basolateral domains: they have a bottle-like shape (Figure 5F).

Several signs suggest that morphological apical-basal polarity in the epithelizing external cells becomes more pronounced. It appears in the position of nuclei, as well as the distribution of septate junctions and elongated granules. Nuclei shift to the sub-apical area (Figure 5E). Septate junctions are mostly located in the subapical regions of external cells initially (Figure 5G), and elongated granules start to concentrate mainly in the apical portion of the cortical cytoplasm (Figure 5H). However, yolk granules and lipid droplets are distributed evenly.

Roundish or polygonal inner non-epithelizing cells are packed more loosely, and well-developed cell junctions between them have never been detected (Figure 5H). Short filopodia-like protrusions cover all surfaces of inner cells, while granules and droplets are distributed evenly in the cytoplasm (Figure 5I).

The number of external cells forming filopodia and lamellipodia (Figure 5J-L: arrows) increases by the end of the early gastrula stage. It was demonstrated in a previous study (Kraus, 2006) that protrusions and crawling activity of external cells provide the merging of epithelial sheet fragments. Thanks to the ultrastructural data obtained, we were able to find out how the formation of these protrusions progresses. At the first step of this process, cells extend their protrusions in all directions (Figure 5J). After that, cells come into contact with their neighbors by using these protrusions (Figure 5K). Cells then reduce almost all protrusions and develop the leading edge (Figure 5L). Finally, cells on the edges of neighboring epithelial sheet fragments contract anchored filopodia and adjacent epithelial sheet fragments composed of up to 20 cells start to merge (Kraus, 2006). With the increase in the area of epithelial sheet fragments and their merging, just several cells of the former flat region end up outside these fragments in the end (Figure 5M, Figure n’).

Epithelial sheet fragments start to bend and their curvature increases during merging. As a result, the flat surface in the center shifts down, and a toroidal surface with a channel in the center forms. The former flat surface turns into the bottom of the torus channel (Figure 5n’’).

### Middle gastrula

At the middle gastrula stage, multiple toroidal surfaces fused with each other compose an embryonic surface. There is a polygonal opening of a channel in the center of each toroidal surface (Figure 6A: arrowheads). Curled free edges of epithelial sheets form inner toroidal surfaces (Figure 5n”, 6B-D).

**Figure 6.**
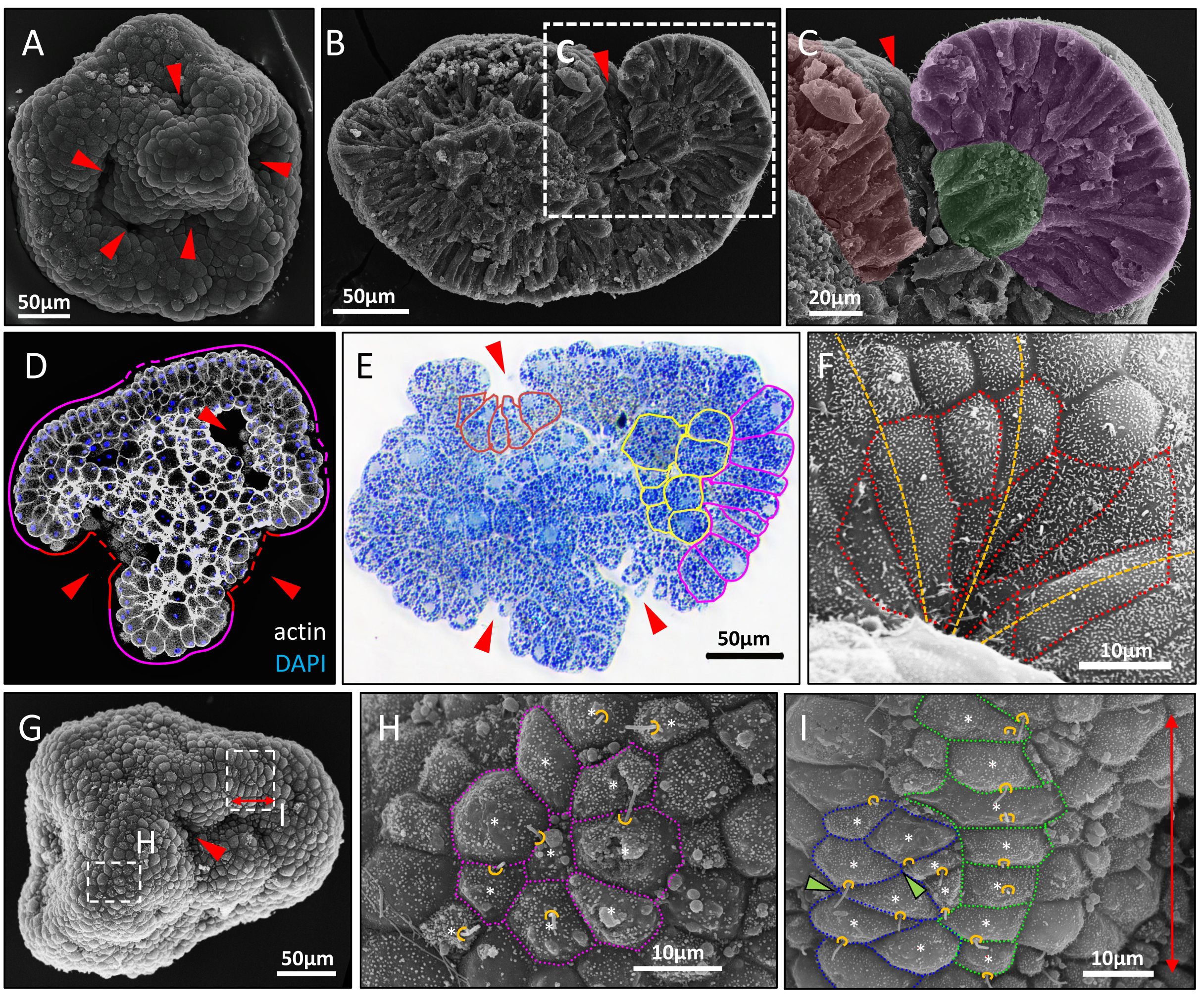
Middle gastrula stage. (A) SEM of the embryo. Red arrowheads indicate torus openings. (B) Middle gastrula split into halves. (C) Rolling up margins of epithelial sheets are colored artificially. (D) CLM central optical section. Magenta lines accentuate epithelialized regions, magenta dotted lines - non-epithelialized regions, red lines - inner toroidal surfaces, red dotted lines - the bottom of the torus channel. (E) Semi-thin section. The shape of cells in the epithelized region is shown by magenta lines, at the bottom of the torus channel - by red lines, inner cells - by yellow lines. (F) The funnel of the torus. Red dotted lines outline cell apices. Yellow dotted curves show meridians of a toroidal surface. (G) Middle/late gastrula. Red double arrow indicates the orientation of the surface fragment on I. (H) Apices of hexagonal cells are outlined in magenta. White asterisks show the geometric center of the cell’s apical surface. Yellow curves show the position of the cilia. (I) Apices of intercalating cells are outlined in blue, apices forming a row - in green. Green arrowheads point to 4-way vertices. SEM: (B, C, F-I).

At the bottom of the torus channel, epithelized cells attain a triangular shape with expanded basal and reduced apical domains (Figure 6E). Non-epithelized cells of the bottom keep a roundish or polygonal shape (Figure 6D). Cells around the opening of a channel have elongated apexes oriented towards the opening (Figure 6F). Cells of outer toroidal surfaces have a columnar shape (Figure 6D, E).

The area of the epithelized surface increases at the middle gastrula stage (Figure 6G). Cilia appear on the external cells (Figure 6H).

The configuration of a cell network, i.e. the orientation and length of interfaces of apical cell surfaces and the number of cell interfaces forming vertices, varies in different regions of the epithelized area. Cell apices are mostly hexagonal within the epithelized region (although penta- and heptagons are also present) and three interfaces form each vertex at the middle gastrula stage. They are located in random positions relative to the geometric center of the cell’s apical surface within this region (Figure 6H).

Other regions, however, are enriched with tetra- and pentagonal cells with no hexagonal cells. Some cells form four-way vertices with their neighbors. Other cells form distinct rows with cell apices elongated perpendicularly to the row length. It seems that this row has its own polarity, which can be visible by the position of cilia relative to the geometric center of the cell’s apical surface (Figure 6I).

### The late gastrula

At the late gastrula stage, the embryo gradually assumes an ellipsoidal shape (Figure 7A). Epithelial sheets continue merging and torus openings start to close. It has been proposed previously that both of these processes proceed by the same mechanism (Kraus, 2006). We confirmed and clarified the details of this mechanism on the ultrastructural level. Cells at the vertices (Figure 7B, magenta arrowheads) of polygonal torus openings form lamellipodia and filopodia (Figure 7C). We observed short filopodia between two cells of two adjacent margins of epithelial sheet fragments (Figure 7D, E). This may indicate that a contraction of filopodia provides a closure of torus openings in a zipper-like manner starting from each vertex of the opening.

**Figure 7.**
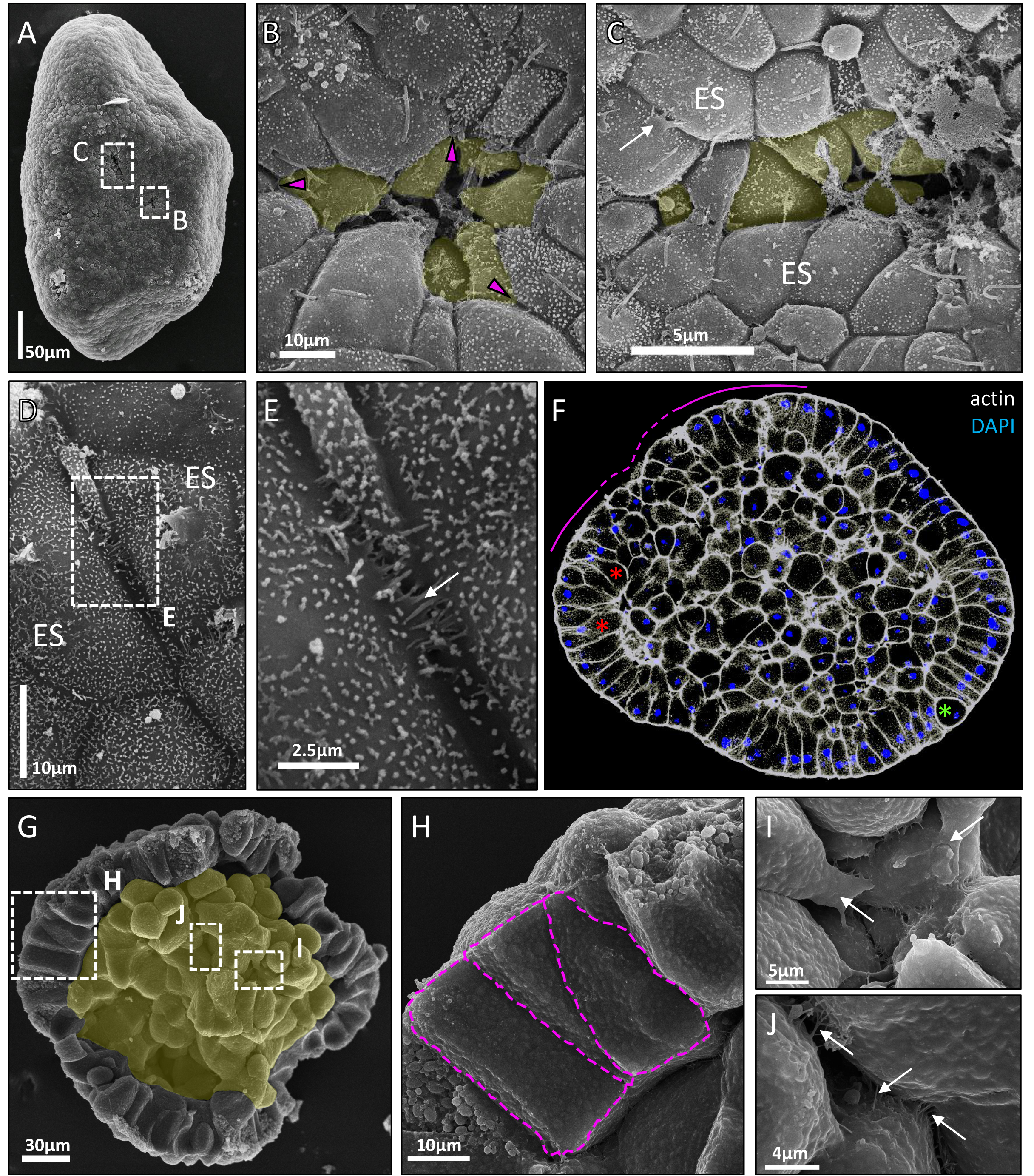
Late gastrula stage. (A) SEM of late gastrula. (B, C) Closure of toroidal openings. Magenta arrowheads point to the vertices of the torus opening. White arrow points to cell protrusions. Deeping inside cells are highlighted in yellow. ES - epithelial sheets. (D) Merging of two ESs. (E) Two adjacent margins of ESs. White arrow indicates short filopodia. (F) CLM of the late gastrula, central optical section. Magenta lines accentuate epithelialized regions, magenta dotted line - non- epithelialized region. Red asterisks - bottle-like cells, green asterisk - rounded cell. (G) Late gastrula split into halves. Unepithelized inner cells are colored in yellow. (H-J) The shape of ectodermal cells (H) is highlighted in magenta. Arrows point to lamellipodia (I) and filopodia (J) of endodermal cells. SEM: (A-E, G-J).

As a result of this zippering, practically the entire embryonic surface becomes epithelized and it is difficult to discern individual toroidal surfaces. In contrast to the epithelized areas, cells do not exhibit a columnar shape in the regions of contact between the edges of epithelial sheet fragments (Figure 7F). It seems that these external cells epithelize later. Cells that form the bottom and the inner surface of the torus come inside the embryo after the closure of the toroidal opening. Later, they join the inner cell mass (Figure 7B, C).

After the closure of the last tori, a continuous superficial epithelial sheet (presumptive ectoderm) is formed (Figure 7G). It consists of columnar and triangular cells (Figure 7H) with single bottle-like cells dispersed among them (Figure 7F: red asterisks). As it normally occurs in the columnar epithelium (Baker, Garrod, 1993; Meyer et al., 2011), mitosis is associated with a surface cell rounding in the ectoderm (Figure 7F: magenta asterisk). The adjacent interphase cells constrict their apical areas, most probably due to compression from rounded mitotic cells according to the data available (Kondo, Hayashi, 2013). The inner cell mass is composed of polygonal cells, which form numerous filopodia and lamellipodia (Figure 7I, J: arrows). One can trace the border separating the external layer of cells and the inner cell mass of the presumptive endoderm (Figure 7G). However, there is still no basal lamina and basal domains of external cells are not well-aligned with each other (Figure 7H).

### The preplanula stage

Gastrulation ends approximately 36-48 hours after the start of development, depending on the temperature. At this stage, embryos of two morphological types can be observed. In the first case, there is an ellipsoid embryo without any opening forms. Oral and aboral poles are poorly distinguishable, but the oral surface is a little rougher (Figure 8A). The ectoderm of the embryo is already completely epithelized and composed of columnar ciliated cells. It surrounds an unepithelized mass of polygonal endodermal cells (Figure 8B). Then a gastric cavity starts to form within the inner cell mass starting from the aboral end (Figure 8C).

**Figure 8.**
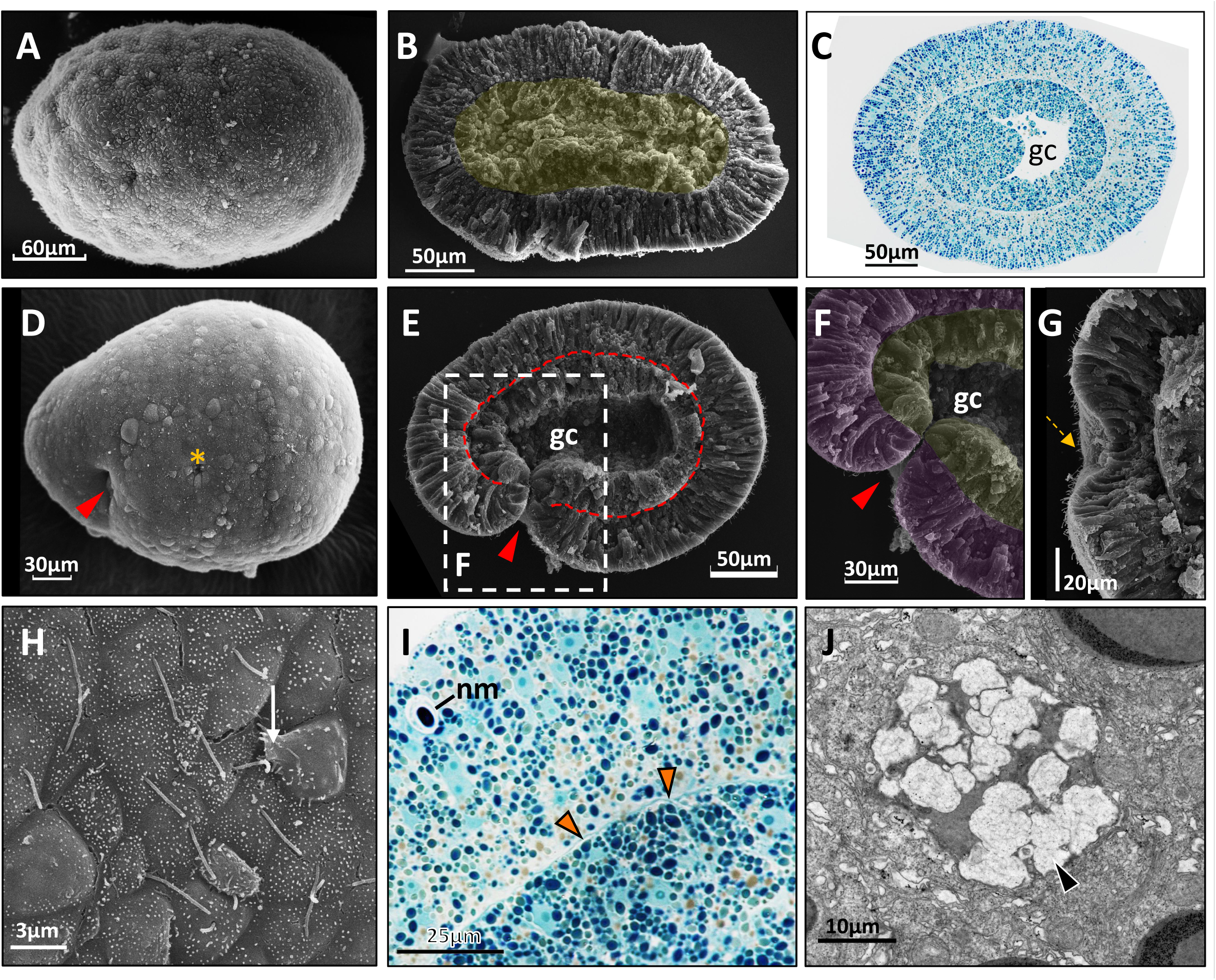
Preplanula stage. The oral pole is on the left at (A, C-E). (A) Preplanula/parenchimula. (B) Parenchimula split into halves. The endoderm is artificially colored in yellow. (C) Semi-thin longitudinal section of preplanula. gc - forming gastric cavity. (D) Embryo with the last torus opening (red arrowhead). Yellow asterisk marks a scar from the closed torus opening. (E) A similar embryo split into halves. The red dotted line shows the boundary between ecto- and endoderm. (F) Presumptive ectoderm is artificially colored in magenta, presumptive endoderm - in yellow. Arrowhead points to the canal connecting the gc with the external environment. (G) Preplanula split into halves. Yellow arrow points to the scar from the last torus opening. (H) Lamellapodia (white arrow), formed by the ectodermal cell. (I) Semi-thin section. Thin basal lamina (orange arrowheads). nc - nematocyst. (J) TEM of mucous vesicle (black arrowhead) in the differentiating glandular cell. SEM: (A, B, D-H).

Embryos of the second morphological type have clearly distinguishable poles and a forming gastric cavity (Figure 8C). The last torus opening is not closed in such an embryo. It is shifted towards the pointed oral (posterior) pole (Figure 8D). A narrow canal connects the forming gastric cavity with the external environment (Figure 8E, F: arrowhead). After the closure of the last opening, ectoderm and endoderm form a continuous layer in the preplanula and only a scar from the last torus opening can be distinguished on the surface of the preplanula near its oral pole (Figure 8G: arrow).

Ectodermal cells of the oralmost area of the embryo form lamellipodia, which can be considered as a sign of cellular motility (Figure 8H: arrow).

Towards the end of this stage, a thin basal lamina appears (Figure 8I: arrowheads). At the preplanula stage, cell differentiation begins. Differentiating nematocytes are clearly distinguishable (Figure 8I). Also in the ectoderm, one can detect cells differentiating into putative mucus-producing glandular cells (Figure 8J).

### The planula larva

The embryonic development of *D. pumila* lasts 80-100 hours (depending on temperature) and ends with the formation of a motile planula larva. In general, the structure of a planula in *D. pumila* is similar to that of other hydroids. A planula swims with a rounded aboral end forward and settles on it during metamorphosis. Its posterior pointed end corresponds to the oral region of the animal. The free-swimming planula gradually elongates (Figure 9A-C).

**Figure 9.**
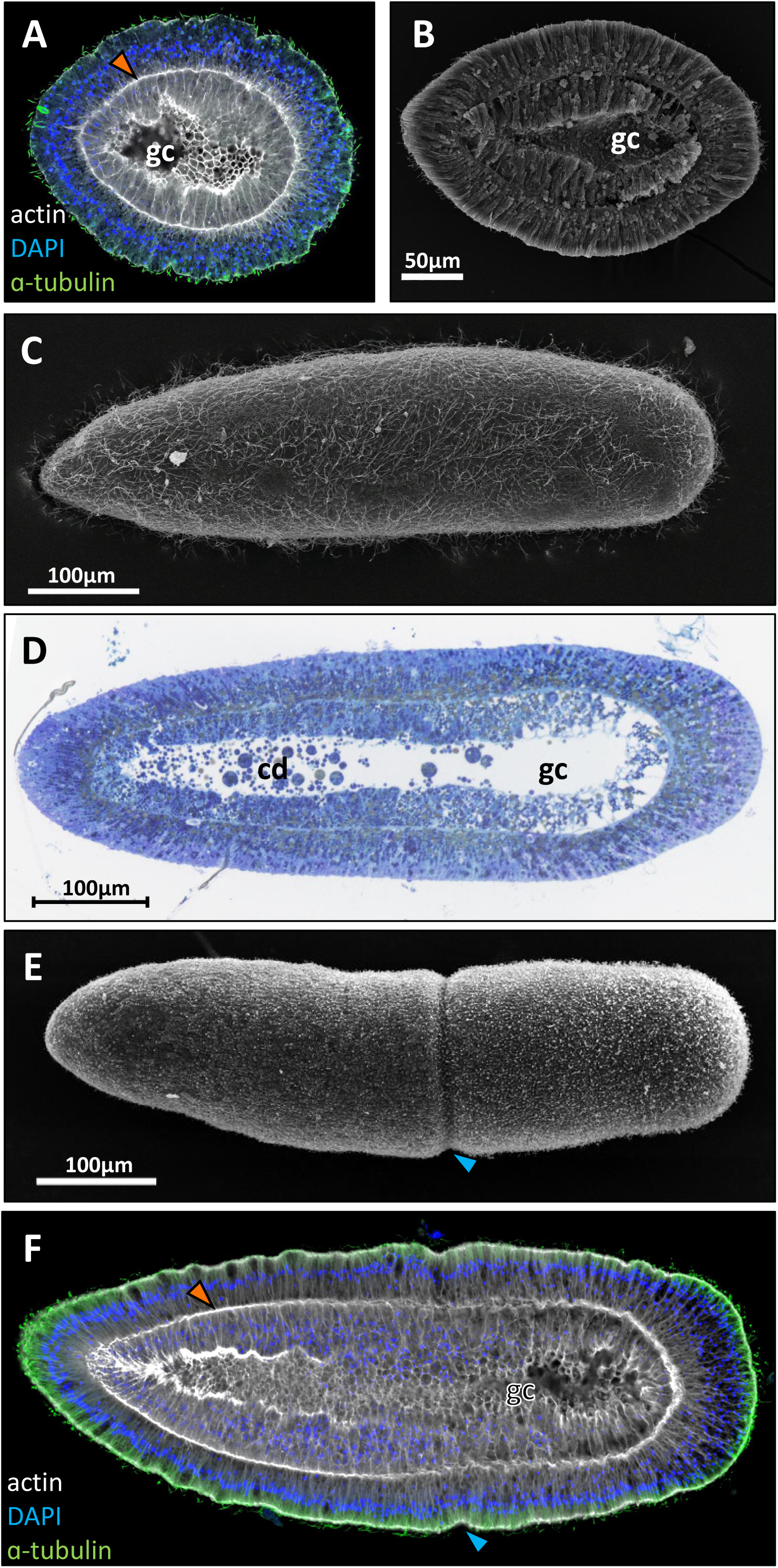
Planula larva. The oral pole is on the left. (A) CLM, central optical section of early planula. Orange arrowhead points to the basal lamina. gc - gastric cavity. (B) SEM of early planula split into halves. (C) SEM of mature planula. (D) Semi-thin longitudinal section of mature planula. cd - cell debris in the gc. (E) SEM of the premetamorphic planula. Blue arrowheads point to ectodermal constriction. (F) CLM, central optical longitudinal section of the premetamorphic planula.

Both ectoderm and endoderm of a planula are fully epithelized (Figure 9B, D). The height of the ectodermal cells is clearly uneven in the fully developed larva. It is about 25–30 μm thick along the body and reaches its maximum (∼60 μm) at the aboral pole. Ectoderm and endoderm are separated by a well-developed basal lamina. The structure of the endoderm typically varies in the mature planula and subdivides into oral two-thirds and oral one-third. In the oral region, endodermal cells have a columnar shape and form a continuous dense layer. Towards the aboral pole the endoderm loses its density, cells become more polygonal and vacuolated. In the gastric cavity, we found cell debris containing lipid droplets and yolk granules (Figure 9D).

In the premetamorpic planula, a circular furrow (Figure 9E, F: arrowhead) appears in the aboral ectoderm not far from the middle of the larva. It is the morphological sign of the planula ready for settlement and metamorphosis.

### Embryo adhesion and spreading

To study embryonic development, we fixed embryos in acrocysts. The acrocyst provides an environment for normal development in natural conditions. Embryos isolated from the acrocyst at stages from the morula and till the late gastrula tend to adhere and to spread on any substrate they are placed on almost into a monolayer (Figure 10A). At the top of the spreading embryo, external cells demonstrate irregular rounded or polygonal morphology (Figure 10B), whereas cells in contact with the substrate flatten and stretch out in the direction of spreading (Figure 10C). Cells at the edge of the spreading embryo are organized into a migratory front and extend wide lamellipodia at their contact with the substrate (Figure 10D). We did not observe the formation of prominent toroidal surfaces in spreading embryos.

**Figure 10.**
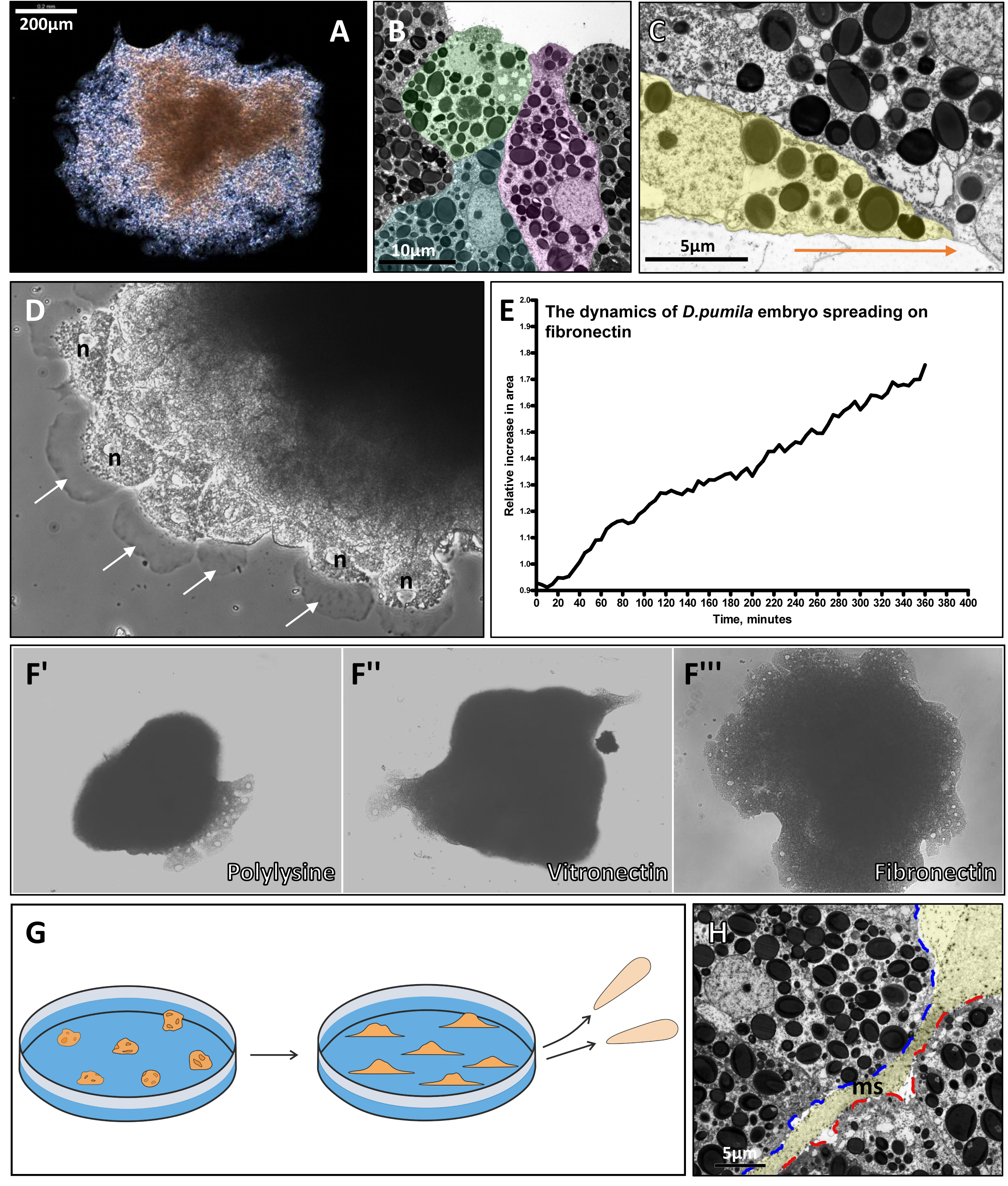
Embryo adhesion and spreading. (A) Embryo spread on plastic during *in vitro* culturing. (B) TEM of cells on the top of the spreading embryo. Artificial coloring accentuates their shapes. (C) TEM of the cell (colored in yellow) crawling over the substrate towards embryo margin. Orange arrow shows the direction of crawling. (D) The morphology of the spreading embryo. Cells form wide lamellipodia (white arrows). n - nuclei. (E) The dynamics of embryo spreading on fibronectin for 6 hours. (F’-F’’’) Gross morphology of embryos after 6 hours spreading on various types of substrates: F’ – polylysine, F’’ – vitronectin, F’’’ – fibronectin. (G) Scheme of the embryo adhesion and spreading. (H) Mucous spacer (ms) between embryos in the acrocyst. Embryos are outlined in red and blue dotted lines. Dence mucus of the spacer is highlighted in yellow.

The spreading of embryos on fibronectin occurs almost linearly without fast or slow phases (Figure 10E). The spreading speed depends on the substrate they are in contact with (Figure 10F’-F’’’). Embryos on polylysine spread weakly, after 6 hours of observation they increase their area 1.21 ± 0.48 fold (N = 3). Similar results were observed in the case of vitronectin (1.15±0.41, N=3). Embryos placed on fibronectin showed the maximum degree of spreading: within 6 hours, the area of embryos increased 1.64 ± 0.28 fold (N = 3), and the cells maintained normal morphology.

Despite such a radical structural reorganization, embryos reduce the area of spreading later on and develop into a normal larva (Figure 10G).

During *in vitro* cultivation, spreading embryos merge sometimes. In natural conditions, adjacent embryos do not stick together, because spacers formed by acrocyst mucous separate them (Figure 10H).

### Cell proliferation in the development of *D. pumila*

To study whether localized regions of cell proliferation exist in the embryo and larva, we used the EdU labeling technique to visualize DNA-synthesizing cells.

In the middle and the late gastrula stage embryos, most EdU-positive nuclei are located in superficial cells (Figure 11A, B).

**Figure 11.**
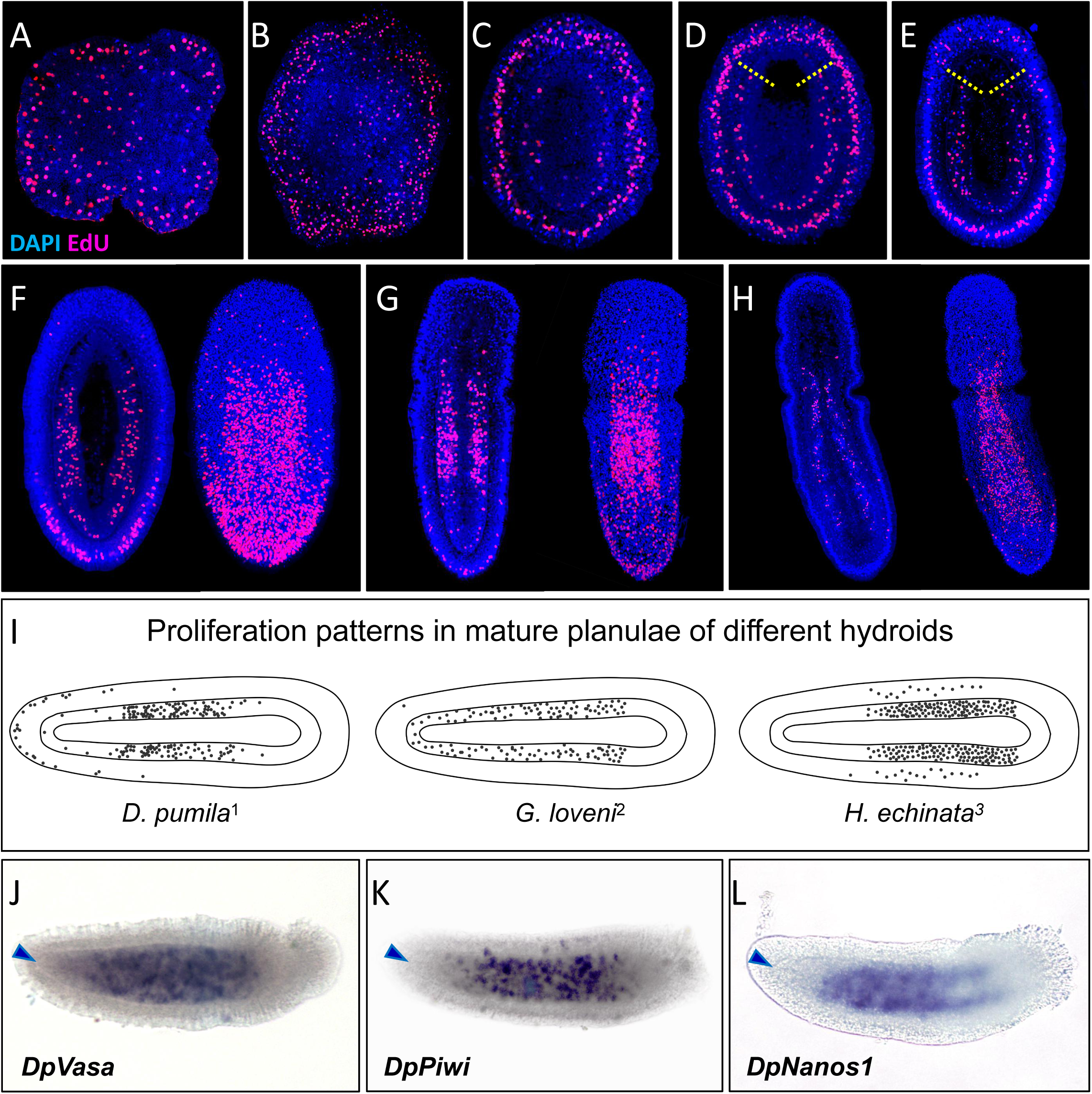
Cell proliferation in the development of *D. pumila.* (A) Middle gastrula. (B) Late gastrula. (C) Parenchymula. (D) Preplanula. Yellow dotted lines show the region of endoderm with low level of cell proliferation. (E) Early planula. (F) Mature planula. (G) Early premetamorphic planula. (H) Late premetamorphic planula. On (F-H) the optical section is on the left, and the 3D reconstruction is on the right. (I) Patterns of cell proliferation in hydrozoan planulae. 1 - *D. pumila*, this study; 2 - *G. loveni*, Burmistrova et al., 2018; 3 - *H. echinata*, Kroiher et al., 1990. (J-K) Expression patterns of DpVasa1, DpPiwi1, DpNanos1. Blue arrowheads point to the border between the ecto- and the endoderm at the oral pole.

At the parenchimula and the preplanula stages, most of the labeled nuclei were found within the ectoderm (Figure 11C, D). In the endoderm of parenchimulae, only a few EdU-labeled nuclei were detected (Figure 11C). In the endoderm of preplanulae, EdU-labeled nuclei are distributed evenly along the oral-aboral axis except the most aboral region and the cell mass in the forming gastric cavity (Figure 11D).

At the early and the mature planula stages, ectodermal cell nuclei predominantly took up the label at the oral end of the larva. In the endoderm, only a few labeled nuclei were found in the aboral-most part of the larva. In the rest of the endoderm, labeled nuclei are distributed evenly (Figure 11E, F).

In the early premetamorphic planula, labeled nuclei are located predominantly in the oral ectoderm. In the endoderm, the EdU-signal was detected mostly in the middle region of the larval body. Only a few labeled nuclei were found in the oral endoderm and only two or three nuclei per optical section could be detected in the aboral region (Figure 11G).

In the late premetamorphic larvae, very few labeled nuclei could be observed in the ectoderm. In the endoderm, we detected no EdU-signal at the poles of the larva, labeled nuclei remain only in the middle region of the larval body (Figure 11H).

Our results suggest that two regions of active proliferation exist in the early premetamorphic larva of *D. pumila*: one is in the endoderm in the middle region of the larval body, another one is in the oral ectoderm (Figure 11G).

One can assume that cell proliferation regions should correspond to the region of multipotent i-cell localization as it was shown for the larva of *H. echinata* (Kanska, Frank, 2013). However, the pattern of cell proliferation in larvae of *D. pumila* differs from that in other hydrozoan species (Figure 11I). That is why this correspondence is not apparent for larvae of *D. pumila.* To clarify the location of i-cells, we analyzed patterns of expression of Piwi, Nanos, and Vasa, established markers for i-cells in hydrozoans (Leclère et al., 2012; Kanska, Frank, 2013; Ruggiero, 2015). *In situ* hybridization revealed DpVasa1, DpPiwi1, and DpNanos1 transcripts in the endoderm in the middle region of the early metamorphic planula (Figure 11J-L). The domain of their expression corresponds to the region of proliferative activity in the endoderm of the early premetamorphic larva (Figure 11G).

### Specification of germ layers in the development of *D. pumila*

To study the specification of germ layers and its dynamics in the embryonic development of *D. pumila*, we visualized the expression of FoxA and Snail, known endodermal markers in hydrozoans (Lapébie et al., 2014; Kraus et al., 2015; Bagaeva et al., 2019; Kraus et al., 2020). For example, their expression is colocalized with an oral region at the beginning of polar gastrulation in *C. hemisphaerica* (Kraus et al., 2020).

In middle gastrula *D. pumila*, *in situ* hybridization with DpFoxA and DpSnail antisense probes revealed their expression in a salt-and-pepper pattern in external and inner cells all over the embryo (Figure 12A, D). Expression of DpFoxA and DpSnail was visualized in the majority of inner cells in the late gastrula and remained in single cells of the external layer (Figure 12B, E). There is no single region of endodermal marker gene expression during the gastrulation in *D. pumila*.

**Figure 12.**
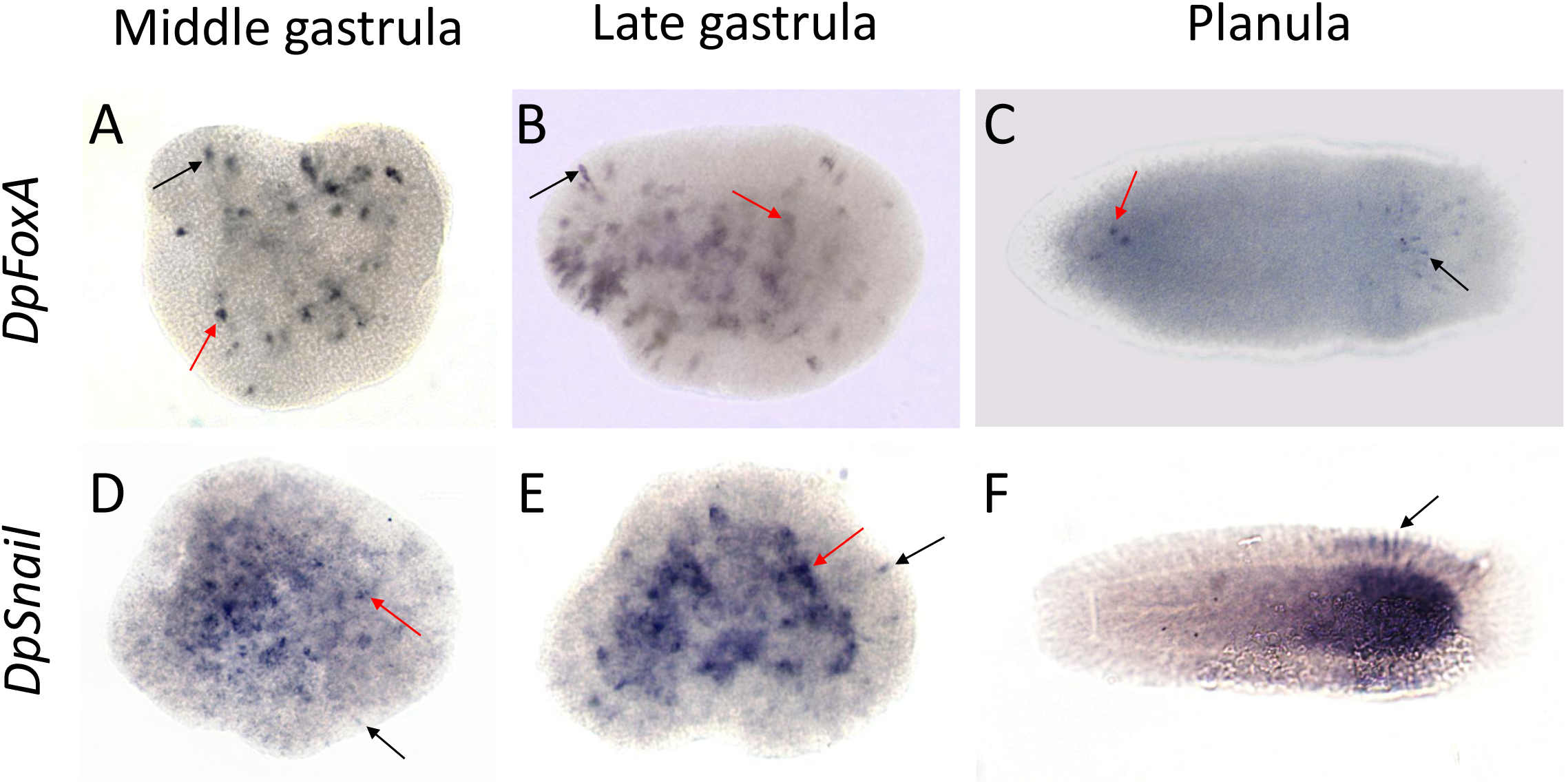
Expression patterns of FoxA and Snail at the stages of middle gastrula, late gastrula, and planula larva in *D. pumila*. Black arrows point to external cells, red arrows - to inner cells, where signal is detected. Oral poles of planulae are to the left.

In the mature planula larva, expression of DpFoxA was visualized only in a few cells scattered along the oral-aboral axis (Figure 12C). A strong DpSnail signal was detected in the aboral endoderm of the larva and the narrow ring of aboral ectodermal cells (Figure 12F).

### The molecular basis of primary body axis formation

It is known that in cnidarians canonical Wnt/β-catenin (cWnt) signaling pathway governs the primary body axis specification and defines a presumptive oral domain in embryos (Wikramanayake et al., 2003; Momose, Schmid, 2006; Plickert et al., 2006; Lee et al., 2007; Momose, Houliston, 2007; Momose et al., 2008; Duffy et al., 2010). In hydrozoans, a key activator of cWnt signaling is Wnt3 (Momose, Houliston, 2007; Momose et al., 2008). In brief, Wnt ligands bind to Frizzled family receptors (Fzd) and activate gene expression, acting through intracellular regulators β-catenin and TCF. When cWnt signaling is not active, destruction complex functions and promotes degradation of β-catenin (Holstein et al., 2011) (Figure 13A). Axin is a core component of the destruction complex and a putative target gene of cWnt (Jho et al., 2002; Kraus et al., 2016).

**Figure 13.**
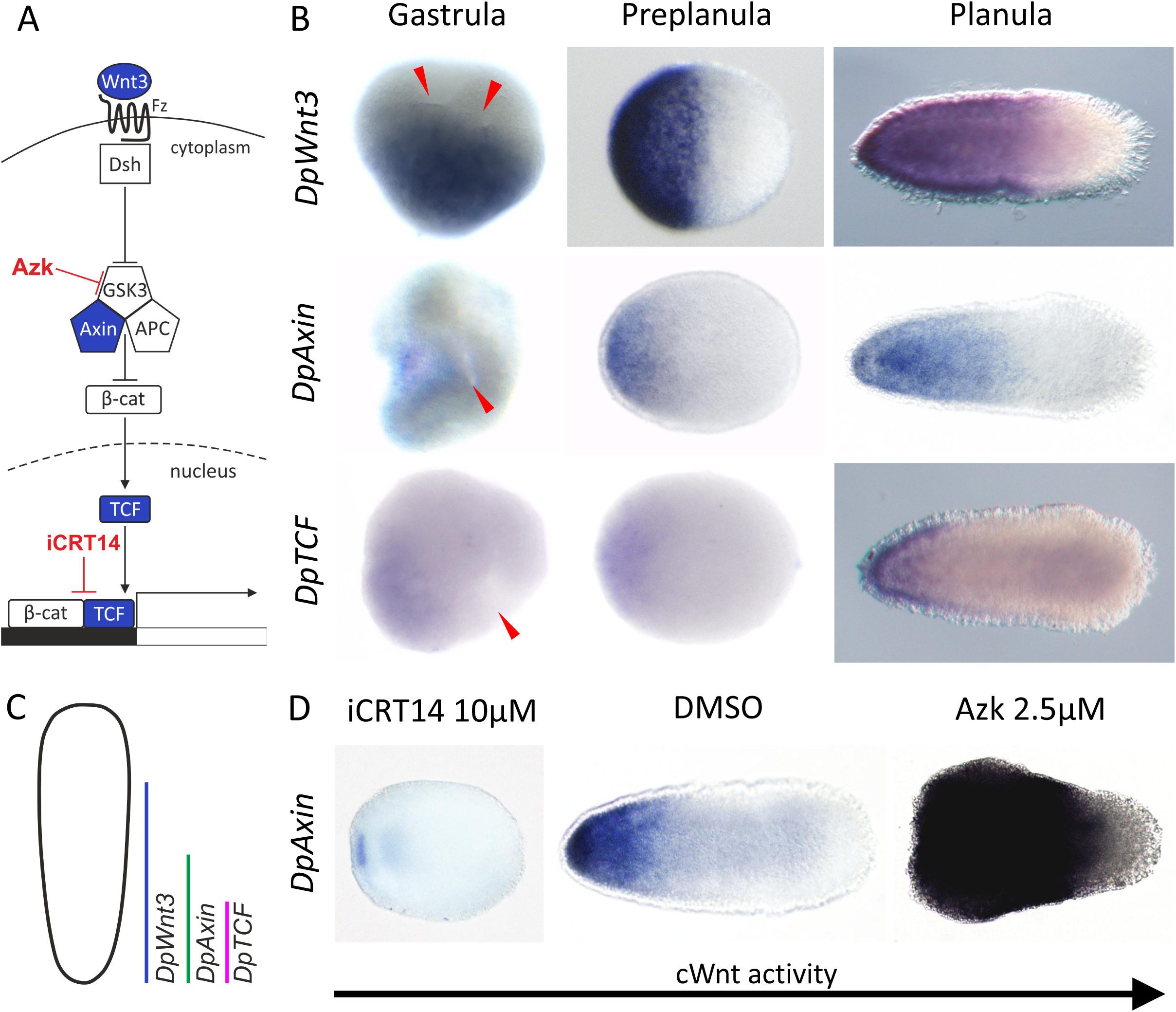
Expression patterns of the cWnt pathway components in the embryonic development of *D. pumila*. (A) The scheme of the cWnt pathway. (B) Expression patterns of the cWnt pathway components at the stages of middle gastrula, preplanula, and planula larva. Red arrowheads point to torus openings. Oral poles of preplanulae and planulae are to the left. (C) The scheme of overlapping expression domains of DpWnt3, DpAxin, and DpTCF in the planula larva. (D) Expression patterns of DpAxin in planula larvae under modulations of the cWnt pathway. iCRT- 14/azakenpaullone (azk) inhibits/activates cWnt signaling.

Since it is impossible to distinguish the direction of the primary body axis based on morphology in embryos of *D. pumila* before the preplanula stage, we studied the formation of the embryo polarity visualizing the expression of cWnt signaling components, DpWnt3, DpAxin, and DpTCF, by *in situ* hybridization (Figure 13B).

We detected a broad domain of DpWnt3 expression and smaller domains of DpAxin and DpTCF expression at the middle gastrula stage. DpWnt3, DpAxin, and DpTCF expression does not colocalize with the formation of epithelial tori or any other morphological features (Figure 13B).

At the preplanula stage, we observed DpWnt3 expression in the one half of the embryo (Figure 13B). DpAxin and DpTCF expression was detected at one of the poles of the preplanula with the weakest signal from DpTCF (Figure 13B).

By the planula stage, DpWnt3 expression covers approximately two-thirds of the larva at the oral end, while DpAxin signal was detected at the oral half of the larva. Signals were detected both in the ecto- and the endoderm. We detected a strong

DpTCF signal in the larval oral ectoderm and dots of weaker expression in the oral endoderm (Figure 13B). Figure 13C visualizes overlapping expression domains of DpWnt3, DpAxin, and DpTCF in the planula larva.

cWnt signaling prepatterns the embryo even when signs of morphological polarity are indistinguishable. To test how cWnt signaling activity affects the formation of the morphological body axis, we treated gastrulae of *D. pumila* with pharmacological modulators of the cWnt pathway (Figure 13A). iCRT-14 blocks the binding of TCF and β-catenin and inhibits cWnt signaling (Gonsalves et al., 2011). Azakenpaullone (Azk) activates cWnt signaling by blocking GSK-3β, a core component of the destruction complex (Kunick et al., 2004; Stukenbrock et al., 2008).

Under inhibition of cWnt signaling with iCRT-14 (10 μM), larvae lack a morphologically distinguishable polarity (Figure 13D). In these larvae, the domain of DpAxin expression was reduced significantly into small patches of stained ectoderm and endoderm. DMSO-treated (control) larvae had normal morphology and expression pattern of DpAxin (Figure 13D). Activation of cWnt signaling with Azk (2.5 μM) resulted in the formation of larvae with an expanded oral domain (Figure 13D). Expression of DpAxin was detected in the entire larva except the aboral-most end.

## Discussion

In our study, we followed the embryonic development of *D. pumila* from the first cleavage division to the planula larva formation (Figure 14). *D. pumila* inhabits a littoral zone. Temperature variations imposed by the ebb and flow of the tide severely affect the rate of development in this species. For example, the rise in temperature by 4°С accelerates the embryonic development of *D. pumila* by 20 hours. Thus, we decided not to create a ‘classical’ table of normal development and focused on embryo morphology to identify and characterize the stages of normal development in *D. pumila*.

**Figure 14.**
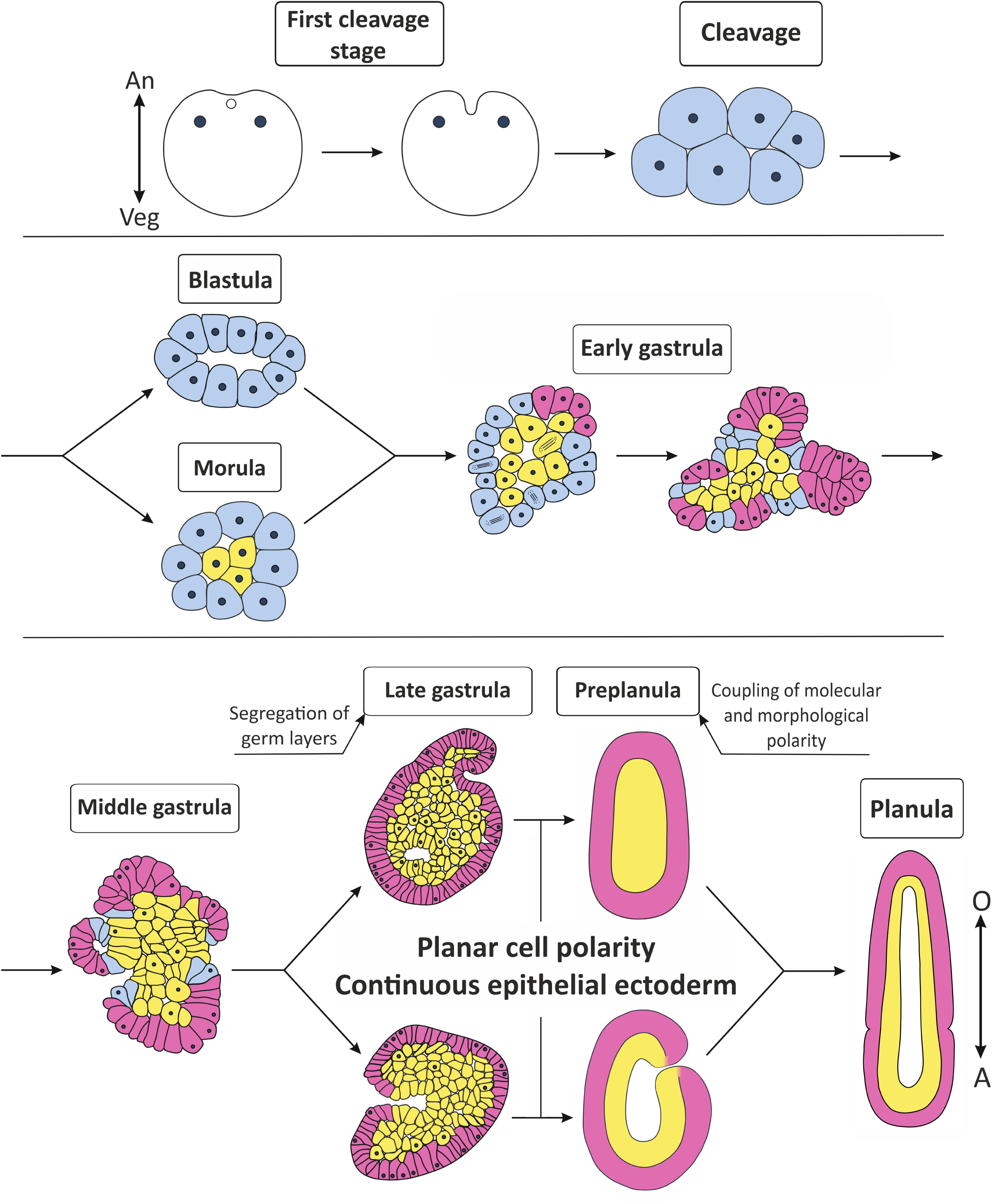
Schematic representation of *D. pumila* embryonic development. Blue cells potentially can give rise or join either ectoderm or endoderm. Presumptive ectoderm is in red, presumptive endoderm is in yellow. Arrows show axes direction: An - animal, Veg - vegetal, O - oral, A - aboral.

### Cleavage and the blastula/morula stages

The arrangement of blastomeres in *D. pumila* could be considered typical for hydrozoans (reviewed by Ivanova-Kazas, 1995). At the 4-cell stage, it imitates the spiral-like arrangement of blastomeres in higher metazoans (Figure 3J). Previously it was described that hydrozoan blastomeres can use each other as a substrate for movement (Kraus et al., 2014; Burmistrova et al., 2018). It seems the same is true for *D. pumila* (Figure 3H), and active translocations of blastomeres provide their spiral-like arrangement. Movements of blastomeres and high cleavage variability results in a high diversity of embryo structure at the 8-cell stage (Figure 2B).

The direction of mitotic divisions is not necessarily uniform during the transition between the 8- and 16-cell stages in *D. pumila*. The morula forms when mitotic spindles are oriented perpendicularly to the embryo surface and daughter cells get inside the embryo (Figure 4B). The blastula forms when all mitotic spindles are directed in parallel to the embryonic surface (Figure 4A; 14). Cleavage also results in blastula and morula in *G. loveni* (Wulfert, 1902; Burmistrova et al., 2018). However, in *H. echinata*, the blastula always forms at the 16-cell stage (Kraus et al., 2014). Unlike *H. echinata*, the contact surfaces of blastomeres never exceed free surfaces in length in the blastulae of *D. pumila*, thus resembling early blastulae in *C. hemisphaerica* (Kraus et al., 2014; Kraus et al., 2020).

Nevertheless, blastula-type embryos always transform into morula-type embryos due to the cell rearrangement and/or the orientation of cell divisions by the 64-cell stage (Figure 4F, G).

### Gastrulation

We assume that cells which get inside the embryo during morula formation will be part of the endoderm, and in *D. pumila*, gastrulation starts as a primary delamination, like in *Hydractinia echinata* (Bunting, 1894; Kraus et al., 2014). So, gastrulation starts when cleavage results in morula formation or during the transition from the blastula to the morula.

Secondary delamination starts with the formation of several epithelial sheet fragments on the curvatures of the embryonic surface (Figure5A, B; 14). Embryos have an irregular shape during this stage and their morphology varies greatly (Figure 2D). In our opinion, macroscopic deformations of the embryo in the acrocyst are the main source of the morphological variability. Embryos are densely packed inside the acrocyst (Figure 1D) (Teissier, 1923) and constrain the geometry of each other. For example, embryonic surfaces become flat where they are in contact with neighboring embryos. Local peaks of surface curvature are observed where flat surfaces meet. Areas of low curvature form where the embryonic surface is in contact with the mucus of the acrocyst (Figure 10H).

On the microscopic level, the main source of variability is the morphogenesis of local epithelial sheet fragments. It proceeds as a coherent process via collective changes in the shape and axis orientation of cells (Figure 5G, 6B) (Kraus, 2006). Therefore, it is impossible to predict the number, size, and location of epithelial sheet fragments in the embryo at the early gastrula stage. During the next step of gastrulation, epithelial sheet fragments merge into toroidal surfaces (Figure 5M, N). The morphological variability decreases. Only macroscopic deformations determine the embryonic morphology at this stage as the number of flat surfaces of the embryo corresponds to the number of forming toroidal surfaces.

On the contrary, the morphological variability rises during gastrulation in *H. echinata*. It is low at the start of gastrulation, but increases dramatically soon. During the so-called folded gastrula stage it reaches its maximum (Kraus et al., 2014).

Folded gastrulae of *H. echinata* bear multiple irregular folds, concavities, and indentations in the embryo surfaces (Kraus et al., 2014). These structures resemble toroidal surfaces in the middle gastrulae of *D. pumila* (Picture 6A), but they have different origins. In *D. pumila,* we suppose that free edges of forming epithelial sheet fragments transform into curling inside margins as the area of epithelial sheets increases (Figure 5N, 6C, 14). Then, epithelial sheet fragments merge due to active cell motility, and multiple fused toroidal surfaces originate (Figure 5N). In *H. echinata*, irregularities on the embryo surface appear to be a side effect of ectoderm overgrowth and an increase of compression on the surface of the embryo. In the cooperative process, groups of compressed cells acquire a bottle-like shape to reduce the compression (Kremnyov et al., 2012). As a result, folds and grooves cover the surface of the embryo (Kraus et al., 2014). In the gastrulae of *D. pumila*, similar compressed cells with a bottle-like shape can be observed, but they are less common (Figure 5F).

Surface deformations do not always accompany the secondary delamination in hydrozoans. The surface of the embryo remains relatively smooth during gastrulation in *Clava multicornis* and *G. loveni* (Ostroumova, Belousov, 1971; Burmistrova et al., 2018).

Further, the morphological variability decreases in both *D. pumila* and *H. echinata.* In *D. pumila*, openings at the center of toroidal surfaces close, and embryos of two morphological types are formed by the end of gastrulation (Figure 14). In the first case, a fully epithelized ectoderm surrounds a mass of non-epithelized endodermal cells. An embryo of this type can be considered as parenchymula (Figure 8B). Other hydrozoans pass through the parenchymula stage in the course of development as well (Metschnikoff, 1886; Kraus et al., 2014; Burmistrova et al., 2018; Kraus et al., 2020), but this does not always happen during the development of *D. pumila*. Gastrulation might result in the formation of an embryo with a cavity, which is connected with the external environment through the last unclosed toroidal opening (Figure 8D-F). It seems that this connection does not have any functional significance.

Previously external cells of walls and the bottom of the torus opening get inside during tori closure passively (Figure 7B, C). Though we were not able to perform cell-labeling, the structure of a preplanula suggests that these cells may contribute to the endoderm population (Figure 8F). With the closure of the last toroidal openings, the segregation of germ layers is completed. Basal surfaces of the ectodermal and the endodermal cells are leveled, and a thin basal lamina appears in such embryos (Figure 8G, I), so we might refer to their ectoderm and the endoderm as epithelial layers by definition (Tyler, 2003).

### “Wound healing” in the gastrulation of *D. pumila*

The merging of sheets of cells with an epithelial zipper is a common process during wound healing in embryonic tissues and epithelium formation in the embryogenesis of higher Metazoans (Raich et al., 1999; Jacinto et al., 2000; Wood et al., 2002; Martin, Parkhurst, 2004; Li et al., 2013). In all these events, opposing cells of epithelial sheets protrude filopodia at the edges, which interdigitate and make weak adhesions in the points of membrane contact. Subsequently, filopodia shorten, drawing cells together, and mature adherens junctions replace the initial weak adhesions. As a result, epithelial sheets may align and adhere together (Yonemura et al., 1995; Vasioukhin et al., 2000; Jacinto et al., 2001; Matilla, Lappalainen, 2008).

Electron microscopy revealed that adherens junction formation in embryos of *D. pumila* proceeds similarly (Figure 3M, N). Cells start to adhere to each other through thin filopodia, which bear spot-like sites of adhesion (Figure 3M). Then, filopodia interdigitate, and adherens junctions increase in length (Figure 3N). Similar filopodial ‘priming’ mechanism also seems to be used, when epithelial sheet fragments merge and the toroidal openings close (Figure 7B-E). Interestingly, irregularities of surface zipper according to the same mechanism in the embryo of *H. echinata* (Kraus et al., 2014). This feature might be quite common for the embryonic development of Hydrozoa with gastrulation via delamination.

It may be assumed that embryonic cells of *D. pumila* perceive unepithelized regions as well as the openings in the center of the epithelial tori as ’’wounds’’, where the integrity of the epithelial layer is broken and should be restored (see also in Kraus, 2003, 2006). Cell behavior resembles lamellipodia-dependent crawling mode of wound healing in these cases (Figure 5J-L, 7B, C). Apparently, when the embryos adhere to a substrate, cells at the edge of the embryo do not perceive contact with other cells on one side. It seems to be enough to cause a migratory behavior in cells on the leading edge. Embryonic cells extend lamellae and crawl forwards (Figure 10D), as occurs during epithelial repair in adult tissues of *Clytia hemisphaerica* (Kamran et al., 2017). It results in embryos flattened and spread drastically (Figure 10).

The degree of embryo spreading was maximal on integrin-specific substrates (fibronectin and vitronectin) in comparison with a non-specific substrate for integrin-mediated adhesion (polylysine) (Figure 8F’-F’’’). Most likely, embryo spreading is an integrin-dependent process. Integrins were reported to promote cell spreading and migration, accumulating in an activated state along the leading edge (Matilla, Lappalainen, 2008). The difference in the dynamics of spreading between the examined substrates can be explained by the expanded repertoire of binding sites for integrins in fibronectin (Gailit, Clark, 1996).

Binding integrins to fibronectin promotes focal adhesion and focal complexes formation (Bachmann et al., 2019). Within hydrozoans, focal adhesions were found in migrating cells in gastrulae of *C. hemisphaerica* (Kraus et al., 2020). According to our experimental data, we may assume that focal junctions are involved in cell migration during the spreading process in *D. pumila*, though we were not able to identify them reliably by microscopy techniques. It is possible that during normal development, cells produce components of the extracellular matrix and form integrin-dependent junctions in the same way as they do when they use each other as a substrate for crawling.

### Morphological differentiation of the primary body axis: morphological landmarks and elongation of the body axis

In *D. pumila*, the morphological polarity of the embryo is visible during the first cleavage (Figure 14). The animal pole can be distinguished by the location of nuclei and the site of the first cleavage furrow formation (Figure 3B-E, G, M). According to Teissier, the site, where the first cleavage furrow forms, corresponds to the prospective oral pole of *D. pumila* planula larva (Teissier, 1931). However, from the second round of cleavage, the morphological axis of the embryo cannot be identified (Figure 3H). The morphological axis is detectable starting from the late gastrula/preplanula stage.

Within metazoans, the main body axis is generated via a convergent extension, a morphogenetic process that causes an embryo to narrow in width as the forming body axis elongates. Planar cell intercalation is a key mechanism for convergent extension during body axis formation (Shindo, 2018; Paré, Zallen, 2020). During planar intercalation, cells exchange neighbors in the plane of the epithelium by cell crawling and/or junction remodeling (Keller, 2002; Devenport, 2016, Shindo, 2018; Huebner, Wallingford, 2018; Paré, Zallen, 2020). On-going cell intercalation can be identified by the configuration of the cell network (Bertet et al., 2004; Blankenship et al., 2006; Lecuit, Lenne, 2007; Tada, Heisenberg, 2012; Shindo, 2018; Keller, Sutherland, 2020).

In *D. pumila* mid-gastrulae, some regions of the embryonic surface are enriched with tetra- and pentagonal cell apices, which form four-way vertices with their neighbors (Figure 6I). Live imaging studies on different species revealed the consistent association of such transitive cell packing with ongoing planar cell intercalation in both mesenchymal and epithelial tissues (Munro, Odell, 2002; Bertet et al., 2004; Nishimura et al., 2012; Williams et al., 2014; Jewett et al., 2017). Thus, we assume that cells repack locally in gastrulae of *D. pumila*. We suppose that four-cell-vertex intermediates subsequently resolve the vertex and establish new interfaces with novel neighbors. Following the completion of these local topological changes, cells intercalate between each other, and distinct cell rows form (Figure 6I). Thus, we suppose that local planar intercalation starts at the end of the middle gastrula stage in *D. pumila*.

Planar intercalation is governed by the planar polarity of cells (Shindo, 2018). A coordinated alignment of cilia can be used as a morphological sign of the planar polarity (Momose et al., 2012; Walentek et al., 2018). In mid-gastrulae of *D. pumila*, such coordinated alignment was detected only in the regions of putative planar intercalation (Figure 6I). Other regions of the embryonic surface exhibit predominantly hexagonal cell packing (Figure 6H) typical for a stable epithelium without any cell rearrangements (Gibson et al., 2006. Lecuit, Lenne, 2007; Lecuit, 2013). We assume that the planar polarity of cells is local at this stage. That is why cell morphogenetic movements proceed independently in different regions of the embryonic surface.

Convergent extension mediated by planar intercalation is regulated and directed by a complex interaction of molecular cues and mechanical factors (Herrera-Perez, Kasza, 2018). Several proteins were identified to play a global-level directional role in the elongation of embryonic tissues (Ninomiya et al., 2004; Pare et al., 2014: Williams, Solnica-Krezel, 2020). The noncanonical Wnt/PCP signaling network serves as a “molecular compass” that orients local planar cell polarization relative to the positional cues within the tissues (Gray et al., 2011). In its turn, planar cell polarization governs the direction of planar intercalation during convergent extension (Shindo, 2018). For example, the gradient of Wnt ligands has been demonstrated to act as a positional cue that orients planar polarity of cells via the non-canonical Wnt/PCP signaling during vertebrate embryonic development (Gao et al., 2011; Gray et al., 2011; Sokol, 2015; Chu and Sokol, 2016).

In the embryo of *C. hemisphaerica*, the global planar polarity of cells is evident at the early gastrula stage (Momose et al., 2012). Cell polarity orientation along the oral-aboral body axis depends on the expression of CheWnt3 at the oral pole, where it is required to activate cWnt signaling. Morpholino-mediated inhibition of CheWnt3 disrupts the elongation of the larval body axis (Momose et al., 2008; Momose et al., 2012). In *D. pumila,* the DpWnt3 expression domain was detected in the middle gastrula (Figure 13B), but unlike *C. hemisphaerica,* the planar polarization of cells appears only locally at this stage (Figure 6I). However, we suppose that cWnt signaling might play a directional role in the establishment of global planar cell polarity later in the embryonic development of *D. pumila*.

Convergent-extension movements are also directed by and require anisotropies in mechanical tensions (Salbreux et al., 2012; Wang et al., 2020). Mechanical tensions arise during tissue morphogenesis and might modulate planar intercalation (Bertet et al., 2004; Rauzi et al., 2008; Aigouy et al., 2010; Chien et al., 2015). In the Drosophila embryo, for example, invagination of the posterior midgut generates a strong pulling force that promotes convergent extension in a germband (Fernandez-Gonzalez et al., 2009; Collinet et al., 2015; Lye et al., 2015; Yu et al., 2016; Kong et al., 2017).

At the middle gastrula stage in *D. pumila*, merged epithelial sheet fragments form multiple fused toroidal surfaces, which compose the surface of the embryo (Figure 6A). Cell apices demonstrate an elongated shape in the funnel of the torus (Figure 6F). Since cell shape may reflect the field of mechanical stresses in tissues (Nestor-Bergman et al., 2018), we assume that the shaping of each toroidal surface produces a local pulling force. Such a local field of mechanical stresses might direct local planar intercalation in *D. pumila* at the middle gastrula stage (Kraus, Cherdantsev, 2003; Kraus, 2006). As gastrulation proceeds, torus openings gradually close (Figure 7A). It was demonstrated experimentally that the last non-epithelized region of embryonic surface such as the last unclosed torus opening tends to be located in the oral half of the forming planula (Kraus, Cherdantsev, 2003). We suppose that the field of mechanical stresses generated by the last toroidal surface might coordinate a global planar intercalation during morphological axis formation in *D. pumila*. Similarly, planar intercalation of ectodermal cells underlying elongation of the body axis was described in *H. echinata* (Kraus et al., 2014).

In *D. pumila,* the last torus opening is the clearly distinguishable landmark of the morphological polarity. Morphological differentiation of the primary body axis occurs during the late gastrula/preplanula transition. The oral domain of the preplanula is more disordered than the aboral one (Figure 8A, D). In *H. echinata*, similar morphological landmarks are detectable at the preplanula stage. The last irregular region of the surface is located at the oral domain of the embryo (Kraus et al., 2014).

The distribution of proliferating cells also becomes asymmetric at the preplanula stage of *D. pumila* (Figure 11D). From this stage onwards, two regions of cell proliferation were detected. In the endoderm, a region of proliferation corresponds to the domain of DpVasa1, DpPiwi1, and DpNanos1 expression at the late planula stage (Figure 11G, J-L). This implies that larval endoderm is a region of i-cell proliferation in *D. pumila*, since Vasa, Piwi, and Nanos are established hydrozoan marker genes for i-cells (Leclère et al., 2012; Kanska, Frank, 2013; Ruggiero, 2015). In the ectoderm of *D. pumila* larva, proliferating cells do not express i-cell marker genes (Figure 11G, J-L). The ectodermal region of proliferation is located at the oral end of the early-late planula larva (Figure 11E-G). The proliferation of ectodermal cells is likely to contribute to the elongation of the larval body axis together with planar cell intercalation (Kraus, Cherdantsev, 2003; Tada, Heisenberg, 2012). This is consistent with the results of (Kraus, Cherdantsev, 2003) that growth of the pointed oral domain promotes elongation of the forming larva significantly. Proliferation almost stops in the oral ectoderm of premetamorphic larva, since it has reached its full length (Figure 11H).

### cWnt signaling in polarization and axis formation in *D. pumila*

In cnidarian species with a polar mode of gastrulation (*N. vectensis*, *C. hemisphaerica*), the region of cWnt activity specifies larval oral domain and coincides with the region of gastrulation morphogenesis (Lee et al., 2007; Momose et al., 2008) and endoderm specification (Kumburegama et al., 2011; Kraus et al., 2020). It was demonstrated that in the anthozoan *N. vectensis*, primary archenteron invagination is Wnt/PCP-mediated and does not require cWnt signaling. However, cWnt signaling mediates endoderm cell fate specification (Kumburegama et al., 2011). Thus, gastrulation morphogenesis coincides with but is uncoupled from the region of cWnt activity in *N. vectensis*.

In *D. pumila, in situ* analyses revealed broad domains of cWnt component expression at the middle gastrula stage, so the oral domain is already prepatterned in the embryo (Figure 13B). However, embryo morphology and morphogenetic movements, i.e. tori formation, are not colocalized with molecular patterning during gastrulation. Moreover, unlike cWnt component expression, there is not a single region of endodermal marker gene expression during gastrulation of *D. pumila* (Figure 12A, B, D, E). We assume that endoderm cell fate specification is not strictly associated with a region of cWnt activity in *D. pumila*. Thus, germ layer formation and morphogenetic processes do not colocalize and could be uncoupled from a molecular axial polarity during gastrulation in *D. pumila*.

From the preplanula stage on, patterns of the cWnt component expression correspond to the embryonic morphology in *D. pumila*. Expression was detected at the oral domain of the preplanula, which is more disordered (Figure 13B). Like *D. pumila, H. echinata* gastrulates via primary and secondary delamination (Kraus et al., 2014). At the late gastrula stage in *H. echinata*, the oral region of cWnt expression coincides with a more disordered area of the embryonic surface (Kraus et al., 2014). Interestingly, the region with a high cell motility corresponds to the cWnt signaling domains both in *H. echinata* and *D. pumila*. It was shown experimentally that the oral domain provides elongation of the planula when the aboral domain was proved to be morphogenetically passive in *D. pumila* (Kraus, Cherdantsev, 2003). Therefore, cWnt signaling may act as a regulator of morphogenesis during larval axis differentiation of *D. pumila*.

In *D. pumila* and *H. echinata*, upregulation of cWnt activity leads to different results (Figure 15), despite similarities in the mode of gastrulation and expression patterns of cWnt components during embryonic development. In *D. pumila*, upregulation of cWnt signaling results in the formation of a larva with a very broad oral domain (Figure 13D). The body axis remains single despite cWnt modulations. In *H. echinata*, activation of the cWnt pathway induces the formation of a multipolar larva with several oral poles (Plickert et al., 2006). Thus, the robustness of the morphological body axis against modulations in cWnt might vary between cnidarian species. The morphological axis forms late in the development of *D. pumila*, but it is highly robust. Transplantation experiments indirectly confirm late formation of the morphological body axis in *D. pumila*. It is known that the oral end of the planula in *D.pumila* can act as an axial organizer during transplantation to another planula larva (Kraus, 2003). The expression of Wnt3 orthologs is characteristic of the axial organizers in Hydrozoa, i.e. for the apex of the hypostome in *Hydra* or the oral tip of the metamorphosing larva in *H. echinata* (Gee et al., 2010; Duffy et al., 2010; Stumpf et al., 2010). We detected DpWnt3 expression at the oral two-thirds of the planula (Figure 13B). However, transplantation of the oral tip of the planula larva into the gastrula does not cause the formation of the secondary axis (Kraus, 2003). The most likely reason is the fact that by the time the morphological axis starts to form, transplanted tissues are completely fused with donor tissues.

**Figure 15.**
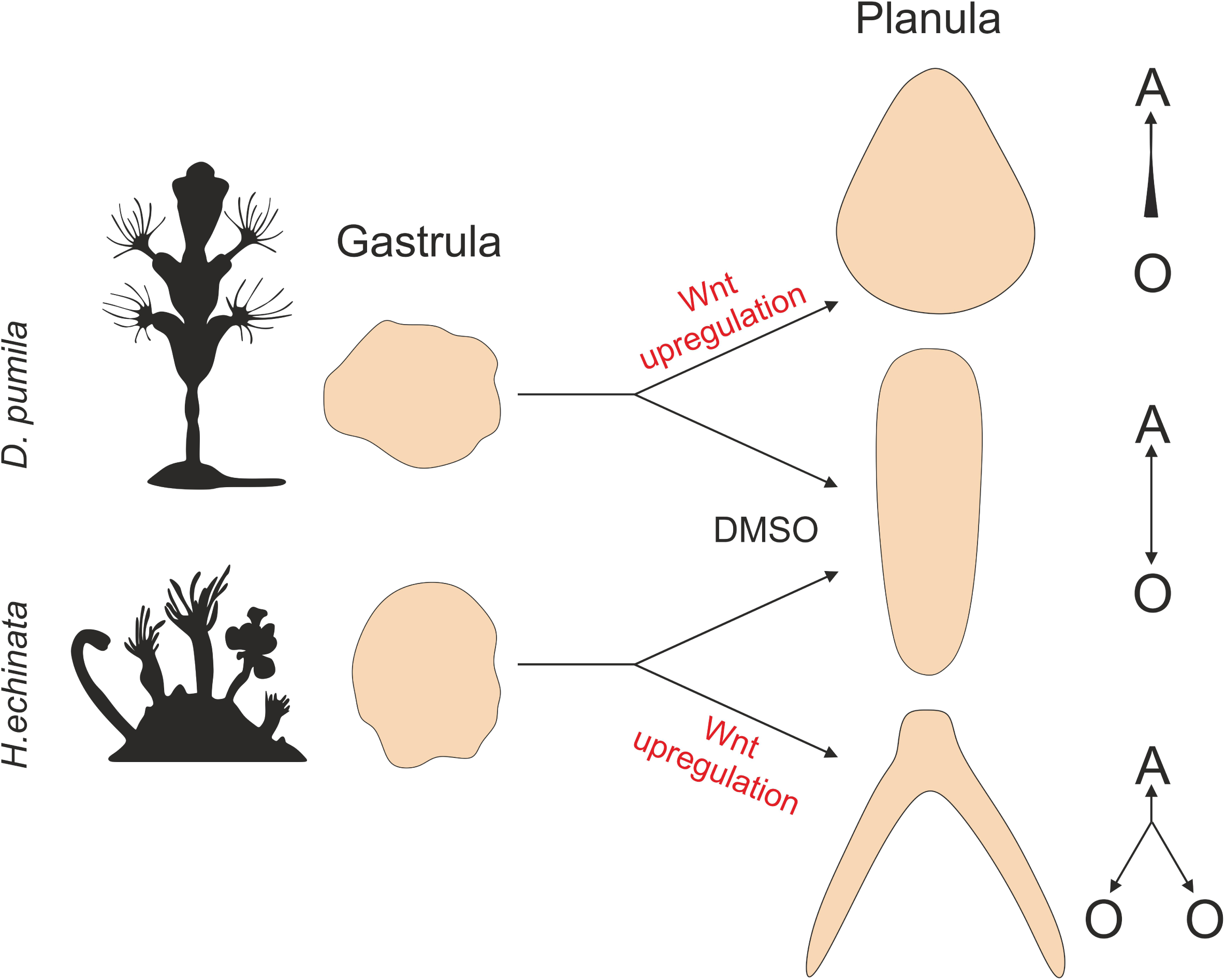
Schematic representation of experiments on upregulation of cWnt pathway in *D. pumila* (this study) and *H. echinata* (Plickert et al., 2006). Arrows show body axes direction: O - oral, A - aboral.

Taken together, our study demonstrates that in *D. pumila,* gastrulation morphogenesis occurs as primary and secondary delamination, in the same way as in several other hydrozoan species. However, in *D. pumila,* gastrulation morphogenesis exhibits unique features and proceeds via the formation of multiple fused toroidal surfaces. Gastrulation morphogenetic processes and cell fate specification are uncoupled from molecular axial polarity based on cWnt signaling. The morphological body axis forms late in the embryonic development of *D. pumila* and is highly robust to modulations in cWnt signaling activity. Further investigation of *D. pumila* might significantly expand our understanding of the links between morphogenesis and axial molecular patterning in Metazoa.

## Experimental Procedures

### Animals

Animal sampling and experimental procedures were performed at the N.A. Pertsov White Sea Biological Station (M.V. Lomonosov Moscow State University) (Kandalaksha Bay; 66°340 N, 33°080 E) during the period of *Dynamena* sexual reproduction (June-July). Sexually mature *D. pumila* colonies were collected and kept in glass bowls filled with seawater at +10-12°C.

### Time-lapse microscopy

We selected embryos at the first cleavage stage and counted the hours after the first division completion. Embryos isolated from the acrocysts and embryos in the acrocysts were placed into Petri dishes filled with filtered seawater (FSW) and kept at +12-16°C. Embryos were photographed every 30 minutes until planula larva formation using the stereomicroscope Leica M165C (Leica, German) coupled to a Leica DFC420 (5.0 MP) digital camera.

### Light and electron microscopy

For light and electron microscopy, specimens were fixed overnight at +4°C in a fixative solution containing 2.5% glutaraldehyde in the buffer solution (0.05 M cacodylate buffer with 15 mg ml^−1^ NaCl (pH=7.2 – 7.4) and 0.025% MgCl_2_). Specimens were then postfixed in 1% osmium tetroxide in the buffer solution at room temperature (2 hours for transmission electron microscopy (TEM), 15 min for scanning electron microscopy (SEM)). They were then washed with the same buffer.

Further processing for histology and TEM was performed as described in Kupaeva et al., 2018. Stained semi-thin sections were examined and photographed under Leica DM2500 (Leica, German) microscope equipped with a Leica DFC420C (5.0MP) digital camera. Contrasted ultrathin sections were examined with the JEM-1011 transmission electron microscope (JEOL, Japan).

Proceedings for SEM were performed as described in Fritzenwanker et al., 2007. Some embryos were split into halves in 70% ethanol. Samples were examined under the CamScan-2 microscope (Cambridge Instruments, UK). Electron microscopy was performed at the Electron Microscopy Laboratory of the Biological Faculty of the Moscow State University.

### Confocal microscopy

Embryos were fixed in 4% paraformaldehyde in FSW overnight at +4°C. Fixed specimens were washed 3×15 min with 1% PBST (1% Triton-X100 in phosphate-buffered saline (PBS)), then treated with 1% PBST for 2 hours.

For immunocytochemical visualization of the mitotic spindles, specimens were treated with a blocking solution (6% bovine serum albumin in PBS) for 2 hours at room temperature and incubated in the primary antibodies against anti-acetylated α-tubulin (mouse monoclonal, 1 mg/ml, Sigma Cat #T7451, USA), diluted in the blocking solution (2 μg/ml) overnight at +4°C. After incubation in primary antibodies, specimens were washed 3×15 with 1% PBST and treated with the blocking solution for 2 hours. Then specimens were incubated with 1.5 μl/ml Goat Anti-Mouse IgG antibodies labeled with DyLight 488 (1.5 mg/ml; Jackson ImmunoResearch Cat #115-486-003, USA) in the blocking solution overnight at +4°C. Samples were then rinsed 3×10 min in PBS. Staining with BODYPI-conjugated phallacidin and DAPI was performed as described in Kupaeva et al., 2018. Optical tissue clearing was achieved by treating embryos with Murray’s Clear solution (2:1 mixture of benzyl benzoate and benzyl alcohol) (von Dassow, 2010). Series of optical sections from samples embedded in Murray’s Clear were obtained with a Nikon A1 confocal microscope (Tokyo, Japan).

### Examination of cell proliferation

For a visualization of cell proliferation, embryos and planulae were incubated in FSW with 20 µM EdU for 2 h at 14-16°C. Then, the samples were washed 3x with filtered seawater and fixed in 4% paraformaldehyde in PBS overnight at 4°C. Further processing was performed according to the manufacturers’ protocol (Click-iT™ EdU Alexa Fluor™ 647 Imaging Kit, #C10340, Thermo Fisher Scientific). Samples were examined with a Nikon A1 confocal microscope (Tokyo, Japan). Z-projections were generated with NIS-Elements D4.50.00 software (Nikon).

### Embryo spreading assay

To perform a qualitative assessment of *D. pumila* embryo spreading on various substrates and a subsequent analysis of its dynamics, we used cover slips coated with 3 types of substrate: polylysine, vitronectin, and fibronectin. To prepare cover slips, they were soaked in concentrated sulfuric acid for 1 hour and then washed in fresh water for the same amount of time. We then placed cover slips into Petri dishes, added 500 μl of substrate solution (20 μg/ml) onto each cover slip, and incubated them for 1 hour at 37°C. After incubation, we let cover slips dry for 1 day at room temperature. Three embryos at the early gastrula stage were placed onto each slip and cultured in FSW. The dynamics of embryo spreading were examined for 6 hours under a Leica M165C (Leica, German) microscope coupled to a Leica DFC420 (5.0 MP) digital camera.

### Chemical treatment

Embryos at the stage of cleavage/early gastrula were treated with 2.5 μM 1-Azakenpaullone (Sigma, Canada/China) or 10 μM iCRT-14 (Sigma, USA/China) to activate/inhibit cWnt signaling respectively. Stock solutions were prepared with DMSO at 10mM, aliquoted and stored at −20°C. To prepare working solutions, stock solutions were diluted in FSW to the final concentration before use. Working solutions were refreshed daily. Control embryos were exposed to 0.1% DMSO in FSW. Incubation was performed in the dark.

### Isolation of *D. pumila* genes, PCR, and antisense RNA probe synthesis

An embryonic cDNA expression library was prepared from total RNA with a Mint cDNA synthesis kit (Evrogen, Russia) using the SMART approach. cDNA fragments of *D. pumila* genes for *in situ* hybridization were isolated from an embryonic cDNA library by PCR with specific primers (see Table 1). Gene-specific primers were designed based upon sequences obtained from the sequenced transcriptome (Illumina) of *D. pumila* (Kupaeva et al., 2020). Orthologous relationships between *D. pumila* genes and previously reported genes were checked via best BLAST hits. Amplified fragments were cloned into the pAL-TA vector (Evrogen, Russia). Digoxygenine- labeled antisense RNA probes were generated from gene fragments, which were transcribed from linearized plasmids containing *D. pumila* genes.

**Table 1:**
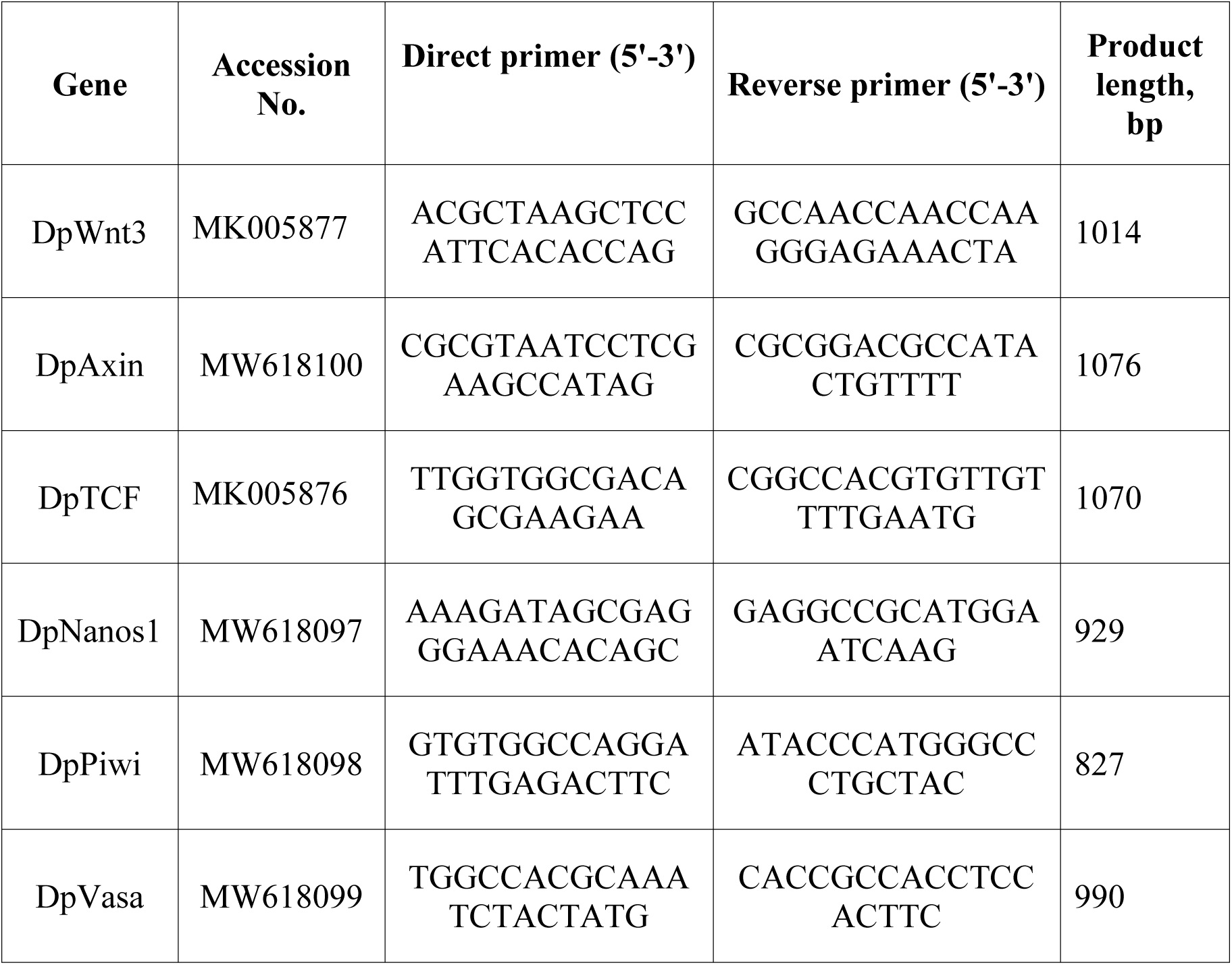
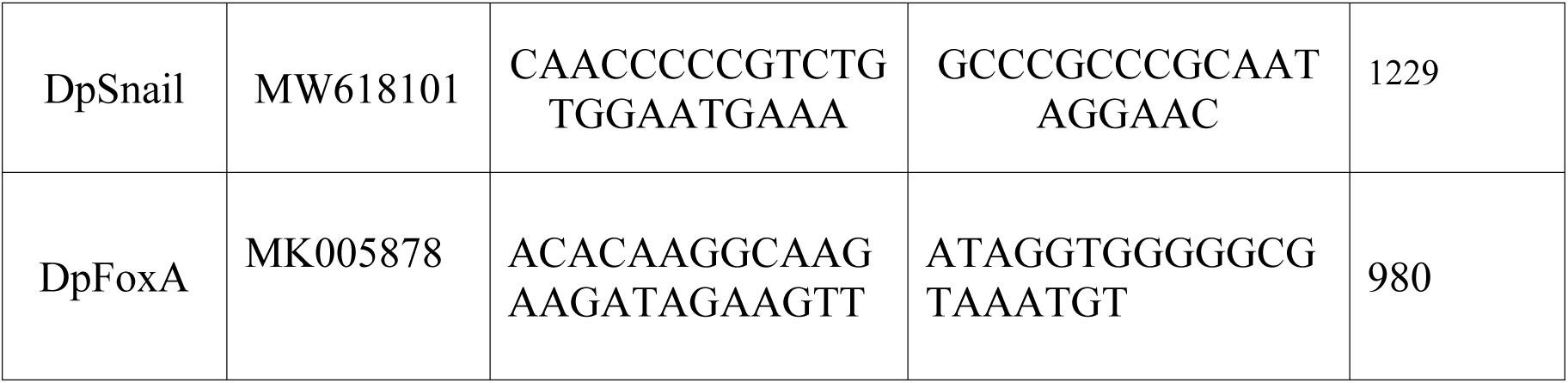
PCR primers for production of ISH probes

### *In situ* hybridization

The *In situ* hybridization protocol was adapted from Genikhovich, Technau, 2009. Embryos were fixed with 4% paraformaldehyde in FSW overnight at 4°C, rinsed with PBS, and stored at -20°C in 100% methanol until hybridization. Samples were rehydrated with PTw (1x PBS with 0.1% Tween 20) and treated with proteinase K (80 μg/ml, 22°C) for 4 minutes. Hybridization was performed at 58°C with digoxigenin-labeled antisense RNA probes (1 ng/μL). Before hybridization, samples were heated at +80°C for 30 minutes for inactivation of endogenous alkaline phosphatase to avoid a false positive result of the reaction. Anti-DIG AP antibody (Roche; 1/2000 diluted) and NBT/BCIP substrate (Roche) were used to detect the probe. After the color reaction, the samples were rinsed with methanol to reduce background staining and mounted in glycerol (87%). Imaging of samples was conducted using a Leica M165C microscope (Leica, German) equipped with a Leica DFC420C (5.0MP) digital camera.

### Image processing

Pictures were edited with ImageJ and Adobe Photoshop CS6 programs. Alterations to the “Brightness’’, “Contrast”, “Exposure”, and “Levels” for the RGB channel were used to achieve optimal exposure and contrast. All tools were applied to the entire image, not locally.

Geometric center of the cell’s apical surface was determined using Icy (v. 2.1.0.1).

## Acknowledgements

We thank the N.A. Pertsov White Sea Biological Station of Moscow State University for the help and support in obtaining samples and providing access to all required facilities and equipment of the “Center of Microscopy WSBS MSU”. We are very grateful to the Shared Facilities center “Electron Microscopy for Life Sciences” of the Lomonosov Moscow State University for the opportunity to perform electron microscopy. We thank N. Grigorian for the language corrections and I. Kosevich for the help and support. The study of embryonic morphology, ultrastructure, and cell behavior during the development of *D. pumila* was funded by RFBR, project number 17-04-01988а. The study of molecular axial patterning during the development of *D. pumila* was funded by RFBR, project number 20-04-00978a. Also, this work was supported by the federal project number 0088-2021-0009 of the Koltzov Institute of Developmental Biology of the Russian Academy of Sciences.

